# Phenotypic determinism and stochasticity in antibody repertoires of clonally expanded plasma cells

**DOI:** 10.1101/2021.07.16.452687

**Authors:** Daniel Neumeier, Alexander Yermanos, Andreas Agrafiotis, Lucia Csepregi, Tasnia Chowdhury, Roy A Ehling, Raphael Kuhn, Raphaël Brisset-Di Roberto, Mariangela Di Tacchio, Renan Antonialli, Dale Starkie, Daniel J Lightwood, Annette Oxenius, Sai T Reddy

## Abstract

The capacity of humoral B cell-mediated immunity to effectively respond to and protect against pathogenic infections is largely driven by the presence of a diverse repertoire of polyclonal antibodies in the serum, which are produced by plasma cells (PCs).^1,2^ Recent studies have started to reveal the balance between deterministic mechanisms and stochasticity of antibody repertoires on a genotypic level (i.e., clonal diversity, somatic hypermutation, germline gene usage).^3–8^ However, it remains unclear if clonal selection and expansion of PCs follows any deterministic rules or is stochastic with regards to phenotypic antibody properties (i.e., antigen-binding, affinity, epitope specificity). Here we report on the in-depth genotypic and phenotypic characterization of clonally expanded PC antibody repertoires following protein immunization. We find that there is only a strong correlation with antigen-specificity among the most expanded clones (top ~ 10), whereas among the rest of the clonal repertoire antigen-specificity is stochastic. Furthermore, we report both on a polyclonal repertoire and clonal lineage level that antibody-antigen binding affinity does not correlate with clonal expansion or somatic hypermutation. Lastly, we provide evidence for convergence towards dominant epitopes despite clonal sequence diversity among the most expanded clones. Our results highlight the extent to which clonal expansion can be ascribed to antigen binding, affinity and epitope specificity and they have implications for the assessment of effective vaccines.

**Graphical abstract:** 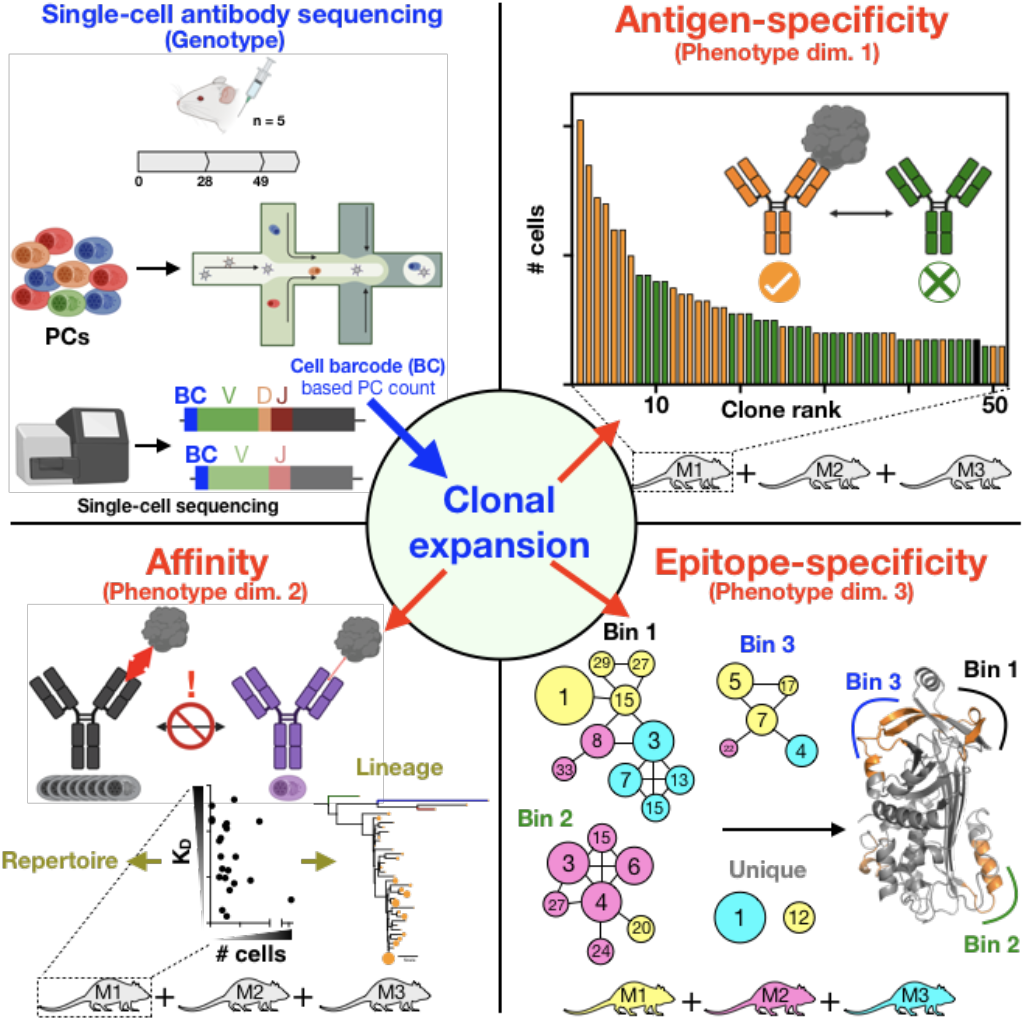

## Introduction

Humoral immunity and successful vaccination require the generation of sustained levels of circulating serum antibodies,^9^ which are produced by clonally expanded PCs, a terminally-differentiated subset of B cells that reside in lymphoid organs (e.g., bone marrow) for an extended period of time (up to years for mice and humans).^1,10,11^ This dynamic process involves the recombination of germline-encoded genetic elements that encode the antibody [or B cell receptor (BCR)] in single B cells;^12^ dogma holds that B cell clonal selection, iterative expansion, and differentiation to PCs occurs for clones with increased affinity towards the antigen.^13–18^ While there have been numerous studies describing how this process is orchestrated on the genotypic level in several species (e.g., humans, mice and zebrafish),^3,4,6,19–21^ much less is known about the associated phenotypic antibody repertoire metrics comprising features such as antigen-binding,^22–25^ quantitative binding affinity and epitope specificity, which can physically be measured as a consequence of the antibody amino acid (a.a.) sequence composition. Importantly, most of the studies reporting phenotypic antibody repertoire data were confined to memory B cells or short-lived plasma blasts that express surface BCR, and in contrast to PCs, do not secrete large amounts of antibody proteins [immunoglobulin (Ig)].^22–24,26–30^

Previous studies on vaccine-induced PC repertoires (murine- and bone marrow-derived) have found that they are dominated by a few (~ 3 - 5) highly expanded clones that are antigen-specific,^19,31^ which correlates with the observation that up to 60 - 90% of the total antigen-specific IgG serum repertoire is comprised of only a few clones (~ 4 - 12).^32–34^ However, it remains unclear whether any deterministic factors such as antigen affinity or epitope specificity drive the selection of these highly expanded clones, and how deep antigen specificity tracks within the PC repertoire. Previous results obtained from adoptive B cell transfer and immunization experiments in BCR transgenic mouse models (monoclonal antibody knock-ins) revealed that high affinity BCRs promote early splenic B cells to differentiate to PCs,^35,36^ a phenomenon that was also reported in a more recent study.^37^ However, contrasting work showed that instead of PC differentiation, the onset of which was later found to occur during late germinal center reactions,^38^ higher BCR affinity led to increased overall proliferation of antigen-reactive cells.^39^ Therefore, it remains unclear how transferable phenotypic antigen binding data from B cells of isolated germinal centers or individual lymph nodes^40,41^ are for the development of long-lived PC repertoires. Here, we set out to comprehensively address these long-standing questions of clonal selection and expansion of PCs. We developed an integrative genotype-phenotype mapping approach and applied it for the in-depth characterization of clonally expanded PC antibody repertoires across five immunized mice.

## Results

We used single-cell sequencing and computational analysis of antibody repertoires combined with quantitative antibody-antigen-binding, -affinity, epitope-binning and -mapping measurements (**Fig. 1a**) which resulted in the characterization of >230 antibodies (which constitute ~ 50% of all captured PCs (IgG) per repertoire). To this end, we repeatedly immunized five BALB/c mice subcutaneously with the T cell-dependent model antigen ovalbumin (OVA) in monophosphoryl lipid A (MPLA) adjuvant. PCs were isolated (based on CD138^+^ B220^low/-^ CD19^low/-^ surface expression) from bone marrow two weeks after the final boost, which represents the peak of PC homing to the bone marrow^42,43^ (**Fig. 1a**). Next, targeted single-cell sequencing was performed on uniquely barcoded heavy and light chain transcripts, covering all Ig isotypes (10X Genomics V(D)J protocol) (**Extended Data Fig. 1a)**. After quality filtering and bioinformatic removal of multiplets, we obtained sequence data on thousands of PCs for each mouse (1891 - 7859 cells) with one paired full-length variable heavy- (V_H_) and variable light- (V_L_) chain sequence (**Extended Data Fig. 1b**), which translated to 672 - 1657 distinct clones (a clone being defined as a unique CDRH3-CDRL3 a.a. sequence) per mouse (across all isotypes) (**Fig. 1b**, **Extended Data Fig. 1c**). To characterize the state of clonal expansion for each repertoire, we operated under the assumption that since the murine naïve BCR sequence diversity (10^13^)^6^ far exceeds the steady-state number of PCs in the bone marrow of a given mouse (~7.5×10^4^),^43^ detecting two or more cells with the same clonal CDRH3-CDRL3 sequence could be used to define clonal expansion (irrespective of isotype origin).^44,45^ We found 199 – 1007 clones to be expanded for each mouse repertoire (30 - 61% of all clones per repertoire), whereas a substantial proportion of clones was detected only once (**Fig. 1b**). Furthermore, all PC repertoires were reproducibly dominated by highly expanded IgM clones (**Fig. 1c, Extended Data Fig. 2**), some of which made up almost 8% of the total cellular repertoire count (IgM-1). As expected, the mutational load between isotypes differed significantly when the 30 most expanded clones per isotype were compared, with most of the IgM clones featuring only a single a.a. intraclonal variant, whereas the number of intraclonal variants markedly increased for expanded IgA and IgG clones (**Fig. 1d, Extended Data Fig. 2b, 3**). Interestingly, when we tested six of the most expanded IgM clones of a single mouse, some of which were also shared between mice, we did not observe any detectable binding to the OVA antigen (or MPLA adjuvant) (**Fig. 1e, Extended Data Fig. 4**). Likewise, when we screened five of the most expanded IgA clones of the same mouse, no antigen-binding was detected (**Fig. 1f, Extended Data Fig. 5**). Therefore, we focused our analysis on the expanded IgG compartment and determined the antigen specificity [by enzyme-linked immunosorbent assay (ELISA)] of the 50 – 60 most expanded antibody clones from mouse (MS) −1, −2 and −5 (which were chosen based on their respective sequencing depth), along with 15 of the most expanded clones from MS-3 and −4 (**Extended Data Fig. 6a, b**). Together, these clones represented 50 - 54% of total IgG cells per mouse for MS-1, −2, and −5 and overall, we found that between 42% and 48% of clones per mouse were antigen-specific (**Extended Data Fig. 6a;** 60% for MS-3 and MS-4). While the most expanded five to ten clones per repertoire were usually antigenspecific, across the rest of the repertoire, antigen-binding and non-binding clones were evenly distributed (**Fig. 1g**). In light of this high degree of stochasticity, we determined how deep antigen-specificity tracks into the repertoire by testing a set of 15 randomly chosen IgG clones from MS-1, including clones that had a cell count as low as one. Even this deep into the repertoire, several clones corresponding to IgG clone rank 249 - 404 and total clone rank 1008 - 1657 showed detectable antigen-binding (**Fig. 1h, Extended Data Fig. 6c, d**).

**Figure 1.**
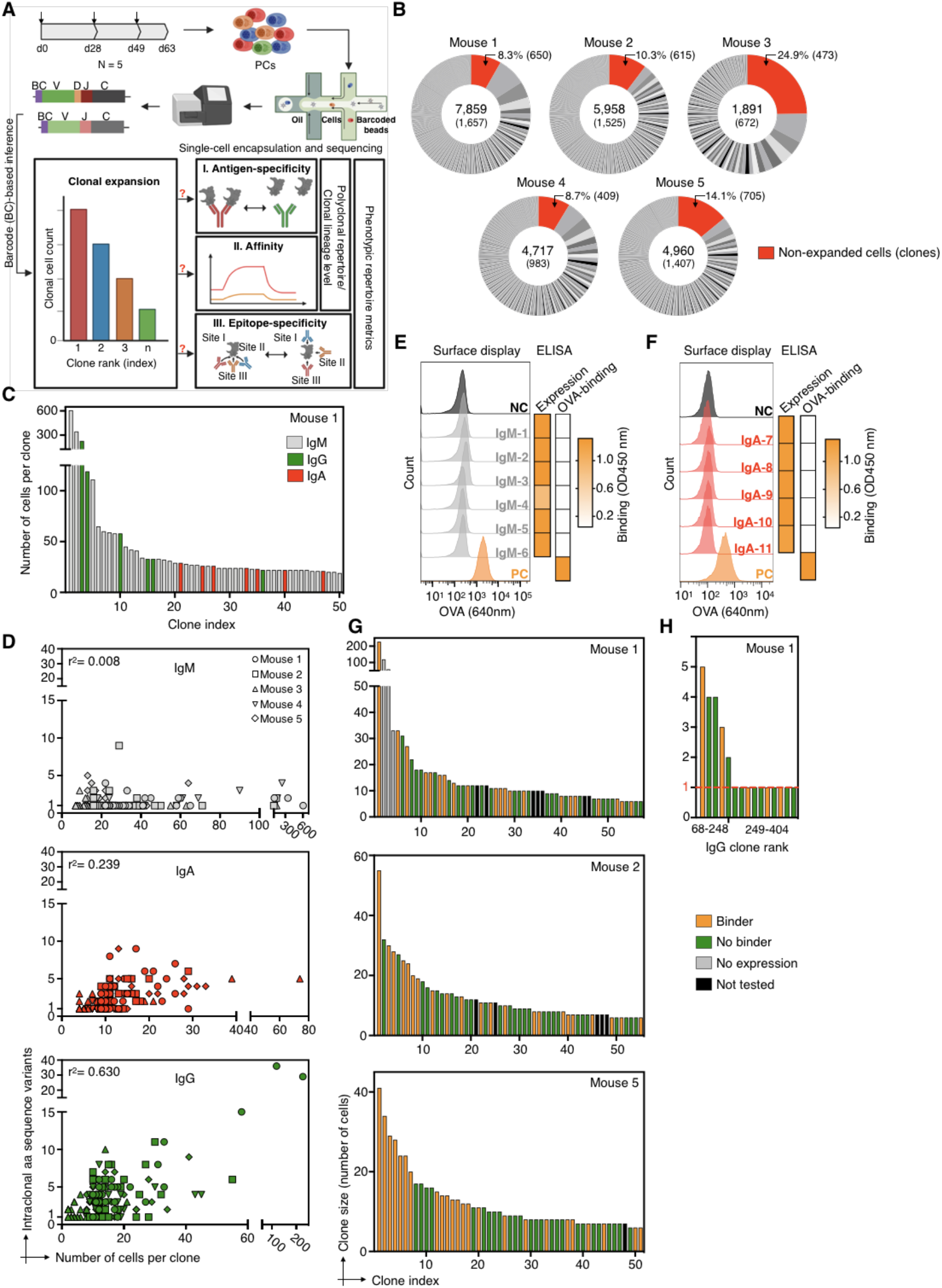
The antigen specificity of clonally expanded plasma cell antibody repertoires. **a.** Schematic project outline. Arrows on top of the timeline graph (top left) indicate time points of mouse immunization. **b.** Pie charts indicate the fraction of productive plasma cells (PCs) per clone and mouse captured by single-cell sequencing (scSeq). Numbers in the center indicate the total number of productive cells (top) and clones (bottom). Unexpanded clones are shown in red and their cellular percentages and total numbers are indicated on top. Clone definition is based on unique CDRH3-CDRL3 amino acid (a.a.) sequence. **c.** Representative clonal expansion profile for the 50 most expanded clones of Mouse 1 (MS-1). Clones are colored by isotype majority and color-coded in grey (IgM), green (IgG) and red (IgA). Profiles of all mice are shown in **Extended Data Figure 2**. **d.** Correlation between the number of intraclonal antibody sequence variants (a.a.) and the number of cells per clone for the 30 most expanded clones per isotype. **e, f.** Screening of six and five expanded IgM and IgA clones of MS-1 for antigen binding. Left: Flow cytometry histogram plot shows ovalbumin (OVA) labeling of hybridoma cell lines with stable surface expression of selected antibody clones (from IgM or IgA PCs) or expression of positive or negative controls (antibodies with defined binding to OVA (PC, orange) or to hen egg lysozyme (NC, dark grey)). Right: heatmaps indicate antibody expression and binding to OVA based on endpoint ELISA (data obtained from **Extended Data Figure 4d and 5b)**. **g.** Antigen-specificity profiling of the top 50-60 expanded clones of MS-1, −2 and −5. Clones with an endpoint ELISA signal >0.2 (three-fold above background) are designated as antigen binders (see also **Extended Data Figure 6a**). **h.** Antigen-specificity of clones showing lower to no clonal expansion (cell count = 1, red dotted line) from MS-1.

Next, given the context of a polyclonal, multi-epitope directed antibody repertoire, we investigated if clonal expansion of PCs is driven by the affinity to the cognate antigen. To this end, we used biolayer interferometry (BLI) to measure the affinity [apparent equilibrium dissociation constants (K_D_)] of 55 antigen-specific IgG clones from MS-1, −2, and −5, resulting in a wide range of affinities (K_D_ = 2 - 450 nM) (**Extended Data Table 1**). This revealed that there was no direct correlation between clonal expansion (based on cell count of a clone) and antigen affinity, as highly expanded clones (cell count > 20) were just as likely to have moderate (K_D_ ~ 100 - 450 nM) or high (K_D_ ~ 10 - 100 nM) affinities, including the surprising observation that the highest affinity (K_D_ < 10 nM) clones often had low cell counts (<10) (**Fig. 2a**). Likewise, we observed highly expanded clones with significantly disparate affinities, as exemplified by the two most expanded clones of MS-5 (5.1 and 5.2), which displayed K_D_ values of 18.7 nM and 451.8 nM, respectively (**Extended Data Table 1**). Moreover, we did not observe a direct link between antibody-antigen affinity and the extent of somatic hypermutation, neither on the a.a. (**Fig. 2b**) nor on the nucleotide (nt) level (**Extended Data Fig. 7**). In some cases, clones that were closer to germline such as the fourth most expanded clone of MS-5 (5.4, 7 a.a. mutations) exhibited high affinity (K_D_ = 8.8 nM), whereas in comparison a highly mutated and highly expanded clone (5.1, 16 a.a. mutations) had a lower affinity (K_D_ = 18.7 nM) (**Extended Data Table 1**). For some antigens, such as influenza hemagglutinin and the spike protein of SARS-CoV-2, it has been shown that certain germlines are structurally predisposed for antigen-specificity and high affinity.^46–48^ Therefore, we determined whether V_H_-V_L_ germline gene (IGHV-IGKV) combinations impacted the affinity of a given antibody to OVA. For the 31 antigen-specific germline combinations tested, some of which shared the same V_H_ or V_L_ genes, we found that affinity was independent of germline gene usage. Most affinities fell into the same range and in case of multiple data points per V_H_-V_L_ combination, the standard deviation for affinity could span a large range (K_D_ ± 250 nM) (**Fig. 2c**). Next, we determined if certain V_H_-V_L_ germline combinations were enriched in the antigen-binding or non-binding fraction of the tested clones. Analysis of V_H_-V_L_ germline gene usage by circos plots of the 79 experimentally verified antigen-binding and 95 non-binding clones did not show enrichment for certain germline combinations in either of the two groups (**Fig. 2d**). Next, we performed a sequence-similarity network analysis (CDRH3-CDRL3 sequence nodes connected by < 4 a.a. difference)^49^ on all IgG clones across all mice (MS-1-5; **Extended Data Fig. 8**) as well as on all experimentally characterized clones. This revealed the presence of discrete subnetworks (**Extended Data Fig. 8, Fig. 2e**), which harbored clusters of both antigen-binding and nonbinding clones, highlighting the potential importance of key amino acid sequence motifs beyond the CDRH3-CDRL3 that contribute to antigen specificity.^50^

**Figure 2.**
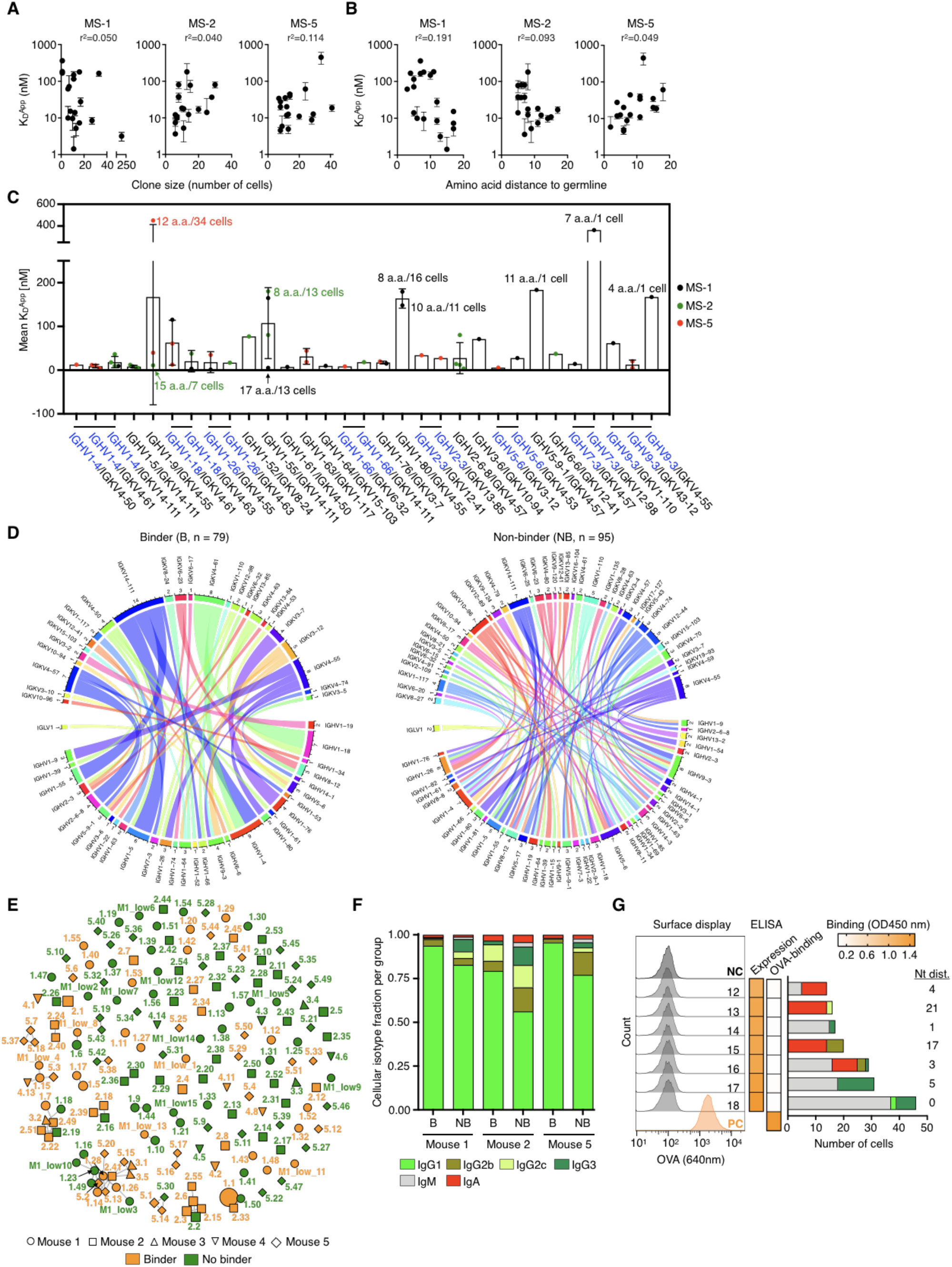
Genotype-phenotype correlations of polyclonal antigen-specific plasma cell repertoires. **a, b.** Correlation between clonal apparent dissociation constant (K_D_) and clone size (number of cells per clone, **a**) as well as clonal a.a. distance to germline (**b**) for MS-1, −2 and −5. Error bars indicate standard deviation (n = 2-3 measurements of K_D_). **c.** Correlation between K_D_ and V_H_-V_L_ germline V-gene usage. V-gene pairs featuring shared V_H_ are indicated in blue with a shared horizontal bar on top. Some data points at the extremes are additionally labeled with the amino acid (a.a.) distance to germline and number of cells for their respective clones. Error bars indicate standard deviation. **d.** Circos plots show the diversity of V_H_-V_L_-gene pairings in the antigen-binding and non-binding group of clones. Each line indicates the germline V-gene usage for an individual VH-VL gene pair. The inner track number and the corresponding thickness of the bar indicate the number of clones utilizing a given germline gene. Color corresponds to the respective germline gene. **e.** Similarity network plot for all 174 antigen-binders and non-binders tested across all mice. Edges represent clones separated by edit distance of < 4 a.a. in CDRH3-CDRL3 sequences. Extent of clonal expansion is reflected by the size of the nodes. Annotated labels of each node are according to clone ID in **Extended Data Table 1** and **2**. **f.** Cellular isotype fractions for all binders (B) and non-binders (NB) per mouse. **g.** Validation of multi-isotype clones by hybridoma surface staining and ELISA screening. Left and middle: flow cytometry histograms and heatmaps similar to as shown in **1e, f**. (ELISA data shown in **Extended Data Figure 11)**. Right: cellular isotype composition of each clone and associated nt distance from germline.

To evaluate whether there were any other features that separate antigen-binding from non-binding clones, we further analyzed common metrics such as distance to germline, number of intraclonal sequence variants, and CDRH3/CDRL3 length and found that only the number of distinct intraclonal antibody sequence variants was significantly increased for antigen-binding clones (**Extended Data Fig. 9**). Upon closer examination of Ig isotype (IgM, IgA) and IgG subtype (IgG1, 2b, 2c, 3), we observed a consistently higher percentage of IgG1 cells among the antigen-binding fraction of clones (**Fig. 2f**). When we tested seven clones that were composed of several isotypes (with minimal to no IgG1), none of them showed detectable antigen binding, suggesting that multi-Ig class-switch recombination does not correlate with antigen specificity (**Fig. 2g, Extended Data Fig. 10**).

Since we established that within the context of the polyclonal repertoire, selection of highly expanded clones was not driven by antigen binding affinity, we next sought to determine the relationship between antigen affinity and clonal expansion within the evolutionary trajectory of a clonal lineage (PCs sharing identical V- and J-genes and identical CDRH3-CDRL3 sequences). We thus expressed antibodies and measured affinities for the most expanded IgG clonal lineage of MS-1, which featured 29 intraclonal sequence variants distributed across a total of 227 cells (**Fig. 3a, Extended Data Fig. 11**). We observed no direct correlation between expansion of an intraclonal sequence variant and binding affinity to antigen. Moreover, the number of somatic hypermutations (distance to germline) did not correlate with affinity and intraclonal variants with lower affinity (K_D_ > 15 - 50 nM) generally originated from earlier ancestral branch points of the lineage tree (**Fig. 3b-d, Extended Data Fig. 12**). This provides further evidence for the existence of a clonal physiological affinity ceiling, as has been proposed in previous work,^24,51^ and which falls within the range of affinities (K_D_ = 2 - 450 nM) measured for polyclonal antibodies (**Fig. 2c**).

**Figure 3.**
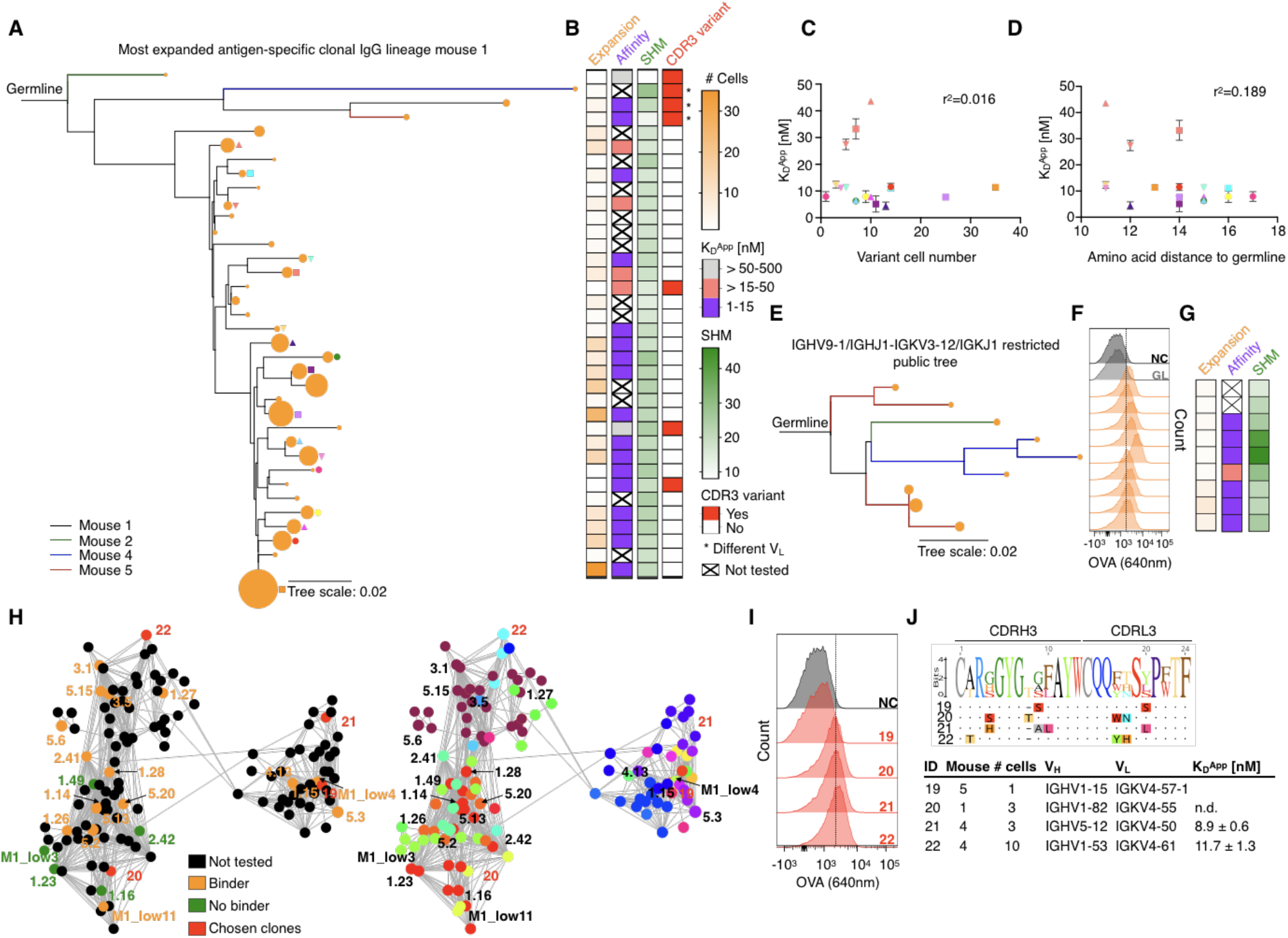
Phenotypic antibody profiling within plasma cell clonal lineages. **a.** Phylogenetic lineage tree of the most expanded IgG clone of MS-1 (see also **Extended Data Fig. 12a** and **b**). Related clones from different mice are indicated by different colors. The size of the orange nodes at the tip of each branch indicates the number of cells per intraclonal variant. Shapes indicate identity of intraclonal variants plotted in **c.** and **d**. **b.** From left to right: heatmaps correspond to intraclonal variant expansion (number of cells per variant), binding affinity (KD), somatic hypermutations (SHM; nucleotide distance to germline) and CDR3 variants (1-6 a.a. edit distance in CDRH3-CDRL3). Clones featuring a different V_L_ are marked by an asterisk. Intraclonal variants from top to bottom correspond to lineage tree variants, as shown in **a. c.** Correlation between apparent dissociation constant (KD) and intraclonal variant cell number for all variants indicated in **a**. Error bars indicate standard deviation (n = 3-5 measurements of K_D_). **d.** Correlation between K_D_ and amino acid distance to germline. Error bars indicate standard deviation (n = 3-5 measurements of K_D_). **e.** Phylogenetic lineage tree of clones originating from several mice that have similar sequences (identical V- and J-genes, CDRH3-CDRL3 with < 5 a.a. difference) (see also **Extended Data Fig. 13a-c**). Branch colors reflect mouse ID from **a.** and node sizes reflect clone size. **f.** Flow cytometry histograms for OVA binding, similar to **1e, 2f.** GL denotes germline, clones correspond to lineage tree shown in **e. g.** Heatmap shown is similar to **3b**. **h.** Network plots of connected IgG sequence nodes from all mice harboring various VH-VL gene combinations. Edges represent clones separated by edit distance of three or less a.a. based on the concatenated CDR3 sequences. Left: verified binders, non-binders, not tested clones as well as newly chosen clones are shown. Clone ID according to **Extended Data Table 1** and **2**. Right: V_H_-V_L_ gene usage visualization. Color code according to **Extended Data Fig. 14c**. **i.** Flow cytometry histograms for OVA binding, similar to **1e, 2f, 3f. j.** Concatenated CDRH3-CDRL3 a.a. sequence logos for clones selected in **h** (red nodes) as well as their antibody characteristics.

Next, we investigated whether clones derived from different mice that shared highly similar sequences (identical V- and J-genes, CDRH3-CDRL3 with < 5 a.a. difference) show similar antigen binding behavior (**Extended Data Fig. 13**). Nine clones were tested, which were similar to a previously identified antigenspecific clone (5.17) and showed consistently similar affinities (K_D_ = 4 - 26 nM) (**Fig. 3e-g; Extended Data Fig. 13**). Next, we examined a major cluster of a sequence similarity network that contained 129 sequence nodes and 32 unique V_H_-V_L_ germline combinations (CDRH3-CDRL3 sequence nodes connected < 4 a.a. difference). Of those, four clones with different germline genes were randomly selected and we could confirm antigen binding for three of them (**Fig. 3h-j**).

Finally, we aimed to understand the epitope specificity and targeting space of expanded PC clones. Using BLI, we first performed cross-competition epitope binning experiments with highly expanded antibody clones that possessed moderate to high affinity (K_D_ ~ 3 - 165 nM). This revealed that within a given mouse, a majority of clones could be grouped into discrete epitope bins (**Fig. 4a, b, Extended Data Fig. 14**). Next, when we tested competitive binding of clones originating from across the different mice (MS-1, −2 and −5) (**Extended Data Fig. 15a**), three dominant epitope bins emerged, some of which contained clones from all mice (bin 1 and 3), whereas others were mainly occupied by clones from a single mouse (bin 2) (**Fig. 4c**). For clones 1.12 and 5.1 (the most expanded clone of MS-5) we did not observe any competitive binding suggesting that they target unique epitopes compared to all other clones tested. Interestingly, we observed a number of different V_H_-V_L_ germline gene combinations for each bin (**Fig. 4c, Extended Data Fig. 15b**). Clonal diversity in each bin was also observed (based on CDRH3-CDRL3 sequences) (**Fig. 4d**) and when we mapped the bin epitope space of the expanded clones on to the complete IgG clonal sequence similarity network (CDRH3-CDRL3 sequence nodes connected < 4 a.a. difference), we could clearly observe epitope convergence despite a high degree of sequence diversity (**Fig. 4e**). This was particularly evident for the biggest sub-network (containing 32 different V_H_-V_L_ combinations) which exclusively harbored clones of epitope bin 1. Phenotypically, we again did not observe a correlation between clone affinity and epitope specificity when affinity data from different mice per bin were compared (**Extended Data Fig. 15c**).

**Figure 4.**
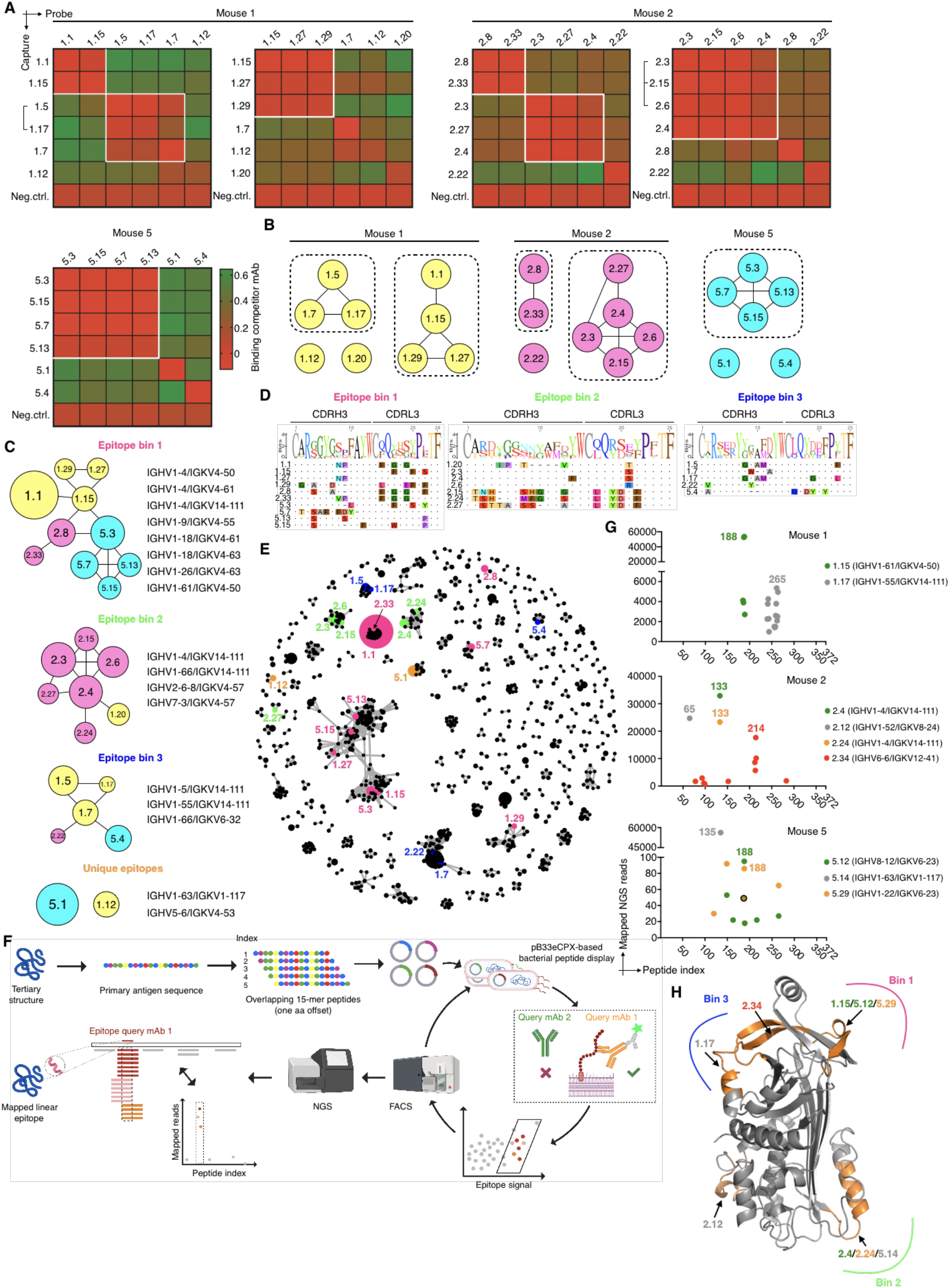
Epitope-targeting space of top expanded clones. **a.** Heatmaps show competitive antigen binding based on BLI assays for highly expanded antibody clones in each mouse. Antibodies indicated on the left were captured and probe antibodies on top were used to determine cross-competition for epitope access. Red indicates no binding of the probe antibody as a consequence of epitope blocking by the capture antibody, whereas green denotes binding of the competitor antibody. Groups of antibodies that target the same epitope (epitope bins) are highlighted in white squares. Brackets indicate clonal variants that share the same VH/VL germline V-genes which differed only in CDRH3/CDRL3 a.a. sequence. An anti-RSVF capture antibody, which does not bind the antigen was used as negative control for all experiments. Clone ID according to **Extended Data Table 1. b.** Epitope bins with associated clones as determined in **a**. Nodes are connected based on observed direct cross-competition. **c.** Epitope bins as defined by the cross-competition of clones from different mice. Representative V-gene combinations are shown on the right. Nodes are connected based on direct cross-competition and sizes indicate clone size (number of cells per clone). Colors represent mouse ID as shown in **b**. Results are reflective of **Extended Data Fig. 16**. **d**. CDRH3/CDRL3 sequence alignment of bin-specific clones. Sequence logo is shown on top and a.a. residues are highlighted if they are in disagreement with the consensus sequence. **e.** Mapping of epitope space as determined in **c** on a sequence similarity network of all IgG clones across all mice (**Extended Data Fig. 9**). Edges represent clones with < 4 a.a. difference in CDRH3-CDRL3 sequence. Node color according to bin color in **c**. Size of clones is reflected by node size. Only those nodes with at least one edge are plotted for visualization purposes. Clone 1.20 is not shown since it was not connected. **f.** Linear epitope-mapping workflow using bacterial peptide display. **g.** Epitope mapping results of select clones from MS-1, −2 and −5. For visualization purposes, only data points with >700 mapped reads are shown for MS-1 and −2 and clone 5.14 of MS-5; for clones 5.12 and 5.29 only data points with >18 mapped reads are shown. Shared data point between 5.12 and 5.29 is indicated with a circle. Corresponding V-gene combinations are indicated. Separate scatter plots featuring all data points are provided in **Extended Data Fig. 18**. **h.** Mapping of epitope bins from **c** on to the OVA crystal structure using antibody epitope information obtained in **g** (PDB: 1OVA).

In order to more precisely define epitope specificity in relation to the sequence and structure of the OVA antigen, we used a bacterial linear peptide display system,^52^ where an overlapping 15-mer (a.a.) peptide library of the primary antigen sequence is expressed on the surface of *E. coli*. Highly expanded antibody clones are then used to label the antigen peptide library and sequential rounds of fluorescence-activated cell sorting (FACS) is performed to enrich for peptide binding, followed by targeted deep sequencing of peptide encoding regions. Subsequently, epitopes are identified using sequence-alignment and read count statistics (**Fig. 4f**). To benchmark the system, we used two commercial monoclonal antibodies with well-defined OVA-specificity for which we could successfully confirm their respective epitopes (**Extended Data Fig. 16**). We then determined the epitopes of nine highly expanded clones across the different mice (MS-1, −2 and −5) (**Fig. 4g, Extended Data Fig. 17**). For all clones (except 2.4 and 5.14, for which only a single highly enriched peptide was found to bind the antibody), FACS enrichment followed by read count analysis resulted in a small number (3 - 12) of overlapping peptide sequences that were highly enriched compared to the background. Within those, we defined the peptide sequence with the highest read count enrichment (observed 86 - 56,000-fold above background) as the corresponding epitope. Importantly, we identified the same epitope for two clones (2.4 and 2.24) sharing highly similar sequences (identical V_H_J_H_-V_L_J_L_ germline, CDRH3-CDRL3 edit distance of 4 a.a.). Finally, several of the clones were also representative of the identified epitope bins (**Fig. 4c**), for example clones 1.15, 1.17 and 2.4/2.24, allowing us to precisely map these bins onto the structure of the OVA antigen, which revealed that they were indeed occupying distinct areas of the antigen (**Fig. 4h**).

## DISCUSSION

Here, we set out to determine if clonal selection and expansion of PC repertoires follows any deterministic rules or is stochastic with regards to their phenotypic antibody properties (i.e., antigen-binding, -affinity, epitope specificity). First, we discovered that except for the most expanded clones (~ top 10), antigen specificity was largely stochastic and could not be predicted based on the number of expanded cells (**Fig. 1g**). Next, we determined if PC repertoire affinities allowed for accommodating the widely accepted germinal center B cell selection model: selective expansion of B cells is driven by an avidity-based selection mechanism in germinal centers involving high-affinity BCRs expressed on B cells that consequentially present the highest levels of peptide-MHC to T follicular helper cells,^41,53,54^ which in turn controls the rate of B cell proliferation.^55^ In contrast to this model, we did not observe any deterministic correlation between antibody affinity and clonal expansion of PCs, furthermore affinity was also not correlated with the number of somatic hypermutations; these stochastic findings were consistent both on the polyclonal repertoire level (**Fig. 2a, b**) and the clonal lineage level (**Fig. 3c, d**).

While we acknowledge that our experimental setup reflects the collective PC-differentiated integration of multiple germinal center outputs that to some extent mirror ongoing (primarily) naïve B cell engagement,^56^ similar results have been observed recently for differentially expanded germinal center B cells.^40^ Therefore, we speculate that over time avidity-based selection may not only be a consequence of affinity alone, but for example can also lead to autoantibody redemption via somatic hypermutation.^57^ Furthermore, from a B cell’s perspective, capturing of antigen may further be modulated by differential BCR stability and surface expression levels, which can also be fine-tuned by somatic hypermutation. To illustrate, recent work has suggested that somatic hypermutation is associated with decreased conformational antibody stability.^58^ Furthermore, highly differential antibody secretion rates from single plasma cells have been reported, which can span up to 3 - 4 log units.^59^ It remains to be determined how these findings translate to surface expression levels of germinal center B cells and subsequent clonal selection, expansion and differentiation to PCs.

Finally, our results also indicate that despite a high level of antibody sequence diversity, the selection of PCs has a deterministic component with regards to specificity towards a few dominant epitopes, which may be attributed to epitope accessibility and availability during B cell maturation and PC differentiation (**Fig. 4c-e**). This could also provide a rationale for the shifting serum epitope immunodominance hierarchies over time, as observed following influenza A virus infection.^60^

Collectively, our findings highlight the fine balance between stochastic and deterministic phenotypic feature selection in differentiated PC repertoires, which only in concert drive clonal selection and expansion. This work has implications for the assessment of vaccine-induced immunity, as understanding how the distribution profile of desired PC clones can be modulated, may support the development of vaccines that provide broad and long-lasting protection by the humoral immune system.

## METHODS

### Mouse immunizations

Mouse immunizations were performed under the guidelines and protocols approved by the Basel-Stadt cantonal veterinary office (Basel-Stadt Kantonales Veterinäramt Tierversuchsbewilligung #2582). Five female BALB/c mice (Janvier Laboratories France, 9 weeks old) were housed under specific pathogen-free conditions and maintained on a standard chow diet. Mice were repeatedly immunized subcutaneously on day 0, 28 and 49 into the flank with 150 μl of a PBS-based immunization cocktail containing 100 μg ovalbumin (Sigma, A5503) and 20 μg monophosphoryl lipid A (MPLA) adjuvant (Sigma, L6895). Blood samples were collected from the tail vein on day 0 and by heart-puncture on day 63. On day 63, mice were euthanized by CO2 asphyxiation and cervical dislocation and femurs and tibias were collected.

### Isolation of plasma cells from bone marrow

Harvested femurs and tibias were clipped with surgical scissors at both ends and the bone marrow was flushed out with ~5 mL of a chilled and sterile filtered solution of PBS pH 7.2, 0.5% BSA and 2 mM EDTA using a 30-gauge BD Micro-Fine+ insulin needle. Bone marrow cells were filtered through a 40 μm nylon cell strainer (FALCON, 352340). PCs defined as CD138^+^ B220^low/-^ CD19^low/-^ antibody-secreting plasma cells were isolated by magnetic activated cell sorting (MACS) using the CD138+ Plasma Cell Isolation Kit mouse (Miltenyi Biotech, 130-092-530) following the manufacturer’s instructions. Briefly, non-plasma cells were first magnetically depleted using a cocktail of biotinylated antibodies against CD49b (DX5) and CD45R (B220) and anti-Biotin microbeads. Then, plasma cells were positively selected from the pre-enriched cell fraction by direct labeling with CD138 microbeads.

### Single-cell sequencing of antibody repertoires

Single-cell sequencing libraries were constructed from the isolated bone marrow plasma cells following the demonstrated 10X Genomics’ protocol: ‘Direct target enrichment - Chromium Single Cell V(D)J Reagent Kits’ (CG000166 REV A). Briefly, single cells were co-encapsulated with gel beads (10X Genomics, 1000006) in droplets using 5 lanes of one Chromium Single Cell A Chip (10X Genomics, 1000009) with a target loading of 13,000 cells per reaction. V(D)J library construction was carried out using the Chromium Single Cell 5’ Library Kit (10X Genomics, 1000006) and the Chromium Single Cell V(D)J Enrichment Kit, Mouse B Cell (10X Genomics, 1000072) according to the manufacturer’s instructions. All of the reverse transcribed cDNA was used as input for VDJ library construction. Final libraries were pooled and sequenced on the Illumina NextSeq 500 platform (mid output, 300 cycles, paired-end reads) using an input concentration of 1.8 pM with 5% PhiX.

### Repertoire analysis

Raw sequencing files arising from multiple Illumina sequencing lanes were merged and supplied as input to the command line program cell ranger on a high-performance cluster. Reads were aligned to the germline using cellranger (v3.1.0) segments from the murine VDJ reference (vdj_GRCm38_alts_ensembl-3.1.0) and subsequently assigned into clonal families based on identical combinations of CDRH3+CDRL3 amino acid sequences. Only those clones containing exactly one productive heavy chain and one productive light chain were retained in the analysis. Isotype majority was determined based on the constant region alignment containing the within each clonal family, with all IgG subtypes merged. Full-length variable sequences (spanning from FR1 to FR4) for each cell were obtained by using the call_MIXCR function in Platypus,^61^ which relies upon aligning full-length contig sequences from the all_contig.fasta file output from cellranger using MiXCR.^62^ To quantify unique variants, full-length VDJRegion (defined as FR1-FR4) for both heavy and light chains were appended together for each cell. Similarity networks were created by calculating the pairwise edit distance using the stringdist package^63^ in R and subsequently creating an adjacency matrix, where each entry (node) corresponded to a unique CDRH3+CDRL3 nucleotide sequence (network shown in **Fig. 2e** was created based on unique CDRH3+CDRL3 a.a. sequence). Edges were drawn between nodes with an edit distance of three or less amino acid mutations and following networks were created by R package igraph. Edit distances were calculated using the stringdist package. Circos plots demonstrating germline gene usage were created using the circlize package^64^ in R. The unmutated germline reference gene was determined by Cellranger’s alignment and set as the outgroup for the phylogenetic tree. Nucleotide and amino acid distance to germline was determined via IMGT.^65^

### Antibody expression

Antibodies were either transiently expressed at small scale in HEK 293 Expi cells using the ExpiFectamine 293 Transfection Kit (Thermo, A14524) and the pFUSE2ss vector system for both, IgM/IgK and IgG/IgK/L (Invitrogen) according to previous protocols,^66^ or stable hybridoma cell lines were engineered by CRISPR/Cas9 genome editing as described before.^67,68^ Of note, these hybridoma cell lines are able to surface display as well as secrete antibody of the IgG2c isotype, which allows for both, FACS-based as well as ELISA-based specificity profiling.

### Hybridoma cell culture

Hybridoma cell lines were cultivated in high-glucose DMEM Medium (Thermo, 61965-026), supplemented with 10% (v/v) of ultra-low IgG FBS (Thermo, 16250078), 100 U/ml Pen/Strep (Thermo, 15140-122), 10 mM HEPES (Thermo, 15630-056) and 50 μM 2-mercaptoethanol (Thermo, 31350-010). Cell lines were maintained at 37 °C, 5% CO2 and passaged every 72 hours.

### Antibody validation by ELISA

0.2 μm sterile-filtered cell culture supernatant of a 6d culture was used to confirm both, antibody expression as well as OVA-specificity. ELISA plates were coated with the capturing reagent in PBS [OVA (Sigma, A5503) for antigen ELISAs, anti-mouse IgM (**γ**-chain specific; Sigma, M8644) and anti-mouse IgG (light chain specific; Jackson ImmunoResearch, 115-005-174) for IgM and IgG expression ELISAs] at 4 ug/ml, blocked with PBS supplemented with 2% (w/v) milk (AppliChem, A0830) and incubated with (serial dilutions of) cell culture supernatant (supernatant of a hen-egg lysozyme/OVA specific cell line served as negative/positive controls respectively). IgM and IgG binding was detected using anti-mouse kappa light chain-HRP (Abcam, ab99617) or anti-mouse IgG (Fc-specific)-HRP (Sigma, A2554) secondary antibody respectively. Binding was quantified using the 1-Step Ultra TMB-ELISA substrate solution (Thermo, 34028) and 1M H2SO4 for reaction termination. Absorbance at 450 nm was recorded on an Infinite 200 PRO (Tecan). All commercial antibodies were used according to manufacturer’s recommendations. For heatmaps shown in **Fig. 1** and **2**, each of the steps was timed to last equally long.

### Antibody validation by surface staining of stable hybridoma cell lines

Flow cytometry scanning of hybridoma cells was performed on a BD FACS Aria III. Typically, 5×10^5^ cells were stained for 30 minutes on ice in 50 μl of a labeling mix consisting of anti-IgG2c-AlexaFluor488 (Jackson ImmunoResearch, 115-545-208), anti-IgK-Brilliant Violet421 (BioLegend, 409511) and OVA-AlexaFluor647 (0.86 mg/ml) at 1:100, 1:80 and 1:50 respectively. Before scanning, cells were washed twice.

### Antibody affinity measurements

Supernatants of Expi cultures and monoclonal hybridoma populations were collected, concentrated (Amicon, UFC810008) and filtered through a 0.2 μm filter (Sartorius, 16534-K). Affinities were then measured on an Octet RED96e machine (FortéBio) with the following parameters: anti-mouse IgG Fc Capture (AMC) biosensors (FortéBio, 18-5088) were hydrated in conditioned media diluted 1 in 2 with kinetics buffer (1xKB) (FortéBio, 18-1105) for at least 10 min before conditioning through 4 cycles of regeneration consisting of 10 s incubation in 10 mM glycine, pH 1.52 and 10 seconds in 1xKB. Conditioned sensors were then loaded with concentrated conditioned medium diluted 1 in 2 with 1xKB (reference sensor) or cell culture supernatant diluted 1 in 2 with 1xKB. Loaded sensors were then equilibrated in 1xKB and typically incubated with various concentrations of OVA antigen in 1xKB ranging from 0 nM (reference sample) up to 200 nM and 1 μM respectively for polyclonal and intraclonal variants mAbs. Finally, sensors were incubated in 1xKB to allow antigen dissociation. Kinetics analysis was performed in analysis software Data Analysis HT v11.0.0.50 and KD-values were calculated from fits with an association R^2^ > 0.85.

### Epitope binning

Transiently expressed antibodies were purified from 30 ml of HEK 293 Expi cultures using Protein G GraviTrap columns (Sigma, GE28-9852-55) and the Ab Buffer Kit (Sigma, GE28-9030-59) according to the manufacturer instructions. Before epitope binning, purified antibodies were confirmed for OVA-positivity by ELISA and OVA-binding kinetics were reconfirmed by BLI measurements prior to each experiment.

Epitope binning following a classical sandwich protocol was performed on an Octet RED96e machine (FortéBio), using anti-mouse IgG Fc Capture (AMC) biosensors (FortéBio, 18-5088) with the following steps: (0) hydration of biosensors in 1xKB for 30 min. (1) Baseline equilibration in 1xKB for 60s. (2) First loading of capture antibody at 40 - 60 μg/ml in 1xKB for 240 s. (3) Quenching of biosensors in 50 μg/ml polyclonal mouse IgG (Rockland, 010-0102) for 300 s. (4) Baseline in 1xKB for 240 s. (5) Second loading of capture antibody at 40 - 60 μg/ml in 1xKB for 240 s. (6) Baseline in 1xKB for 200 s. (7) Loading of OVA at 150 nM in 1xKB for 600 s. (8) Baseline in 1xKB for 60 s. (9) Loading of probe antibody at 25 μg/ml in 1xKB for 600 s. (10) Regeneration of sensors. Analysis was performed in analysis software Data Analysis HT v11.0.0.50.

### Library generation for epitope mapping

Bacterial epitope mapping was based on the pB33eCPX plasmid, which was previously established^52^ to fuse a peptide to the enhanced outer membrane protein X (OmpX) for extracellular peptide presentation. First, we modified the plasmid using primers EpMap_1 and EpMap_2 along with ssODN EpMap1_ssODN (**Extended Data Table 3**) using the NEBuilder Mastermix (NEB, E2621S) following the manufacturer’s instructions in order to create hairpin-free overlap regions that could be used for homology-based library cloning. Next, the epitope library was generated using MC1061 *E. coli* cells, NEBuilder Mastermix, primers EpMap_3 and EpMap_4 (**Extended Data Table 3**) along with 0.078 pmol of a ssODN library of 105 nucleotides in length [30 bp homology overhangs on each side (GGAACTTCTGTAGCTGGACAATCTGGACAA and GGAGGGCAGTCTGGGCAGTCAGGTGATTAC) flanking a 45 bp stretch encoding a 15mer amino acid peptide sequence; ssODN library was ordered as a Tier 1 oligo pool from Twist Bioscience]. The library was designed to cover the whole OVA amino acid sequence, with a window size of 45 bp and a cutoff of one codon, resulting in a library size of 372. The final library size was determined to contain 268,900 transformants using serial dilutions (722x oversampled) and when we analyzed 20 clones by Sanger sequencing using primer EpMap_5, we found that 95% contained a correct sequence of which 95% were unique. Glycerol stocks were subsequently stored at −80 °C until further use.

### Epitope mapping by bacterial surface display

Bacterial display-based epitope mapping was carried out as described before.^69^ Briefly, 500 ml of LB-medium supplemented with chloramphenicol (34 μg/ml) and 0.2% (w/v) 0.2 μm sterile filtered glucose were inoculated with the transformant library and grown for 12 hours at 37 °C. Next, cells were subcultured 1:50 into 5 ml of LB supplemented with chloramphenicol and grown for 2 hours at 37 °C, before protein expression was induced for 1 hour at 37 °C using 0.04% (w/v) L-arabinose. Typically ~1.1E7 cells were subsequently harvested at 3000 g for 5 minutes, washed and stained for 30 minutes on ice in 100 μl of a labeling mix consisting of 20 μl of concentrated and extensively PBS buffer-exchanged hybridoma cell culture supernatant (containing antibody protein of the top expanded clones) and 80 μl of PBS pH 7.2, 0.5% BSA and 2 mM EDTA [for assay establishment, 0.5 μg of control antibodies Clone 4B4E6 (Chondrex, 7096) and Clone 2322 (Chondrex, 7094) were spiked into concentrated and buffer exchanged cell culture supernatant of a hybridoma cell line, that did not express antibody]. Cells were washed twice and stained in 100 μl for 30 minutes on ice using an anti-mouse IgG-Brilliant Violet421 secondary antibody at 1:90 (BioLegend, 405317). Cells were again washed twice, resuspended in 500 μl of PBS and sorted into sterile SOC media using a BD FACS Aria III. Typically, between 10,000 and 50,000 cells were sorted on 2 - 3 consecutive days to enrich for pure antigen-binding populations, before plasmid was extracted and sequencing libraries were generated.

### Generation of epitope mapping sequencing libraries

Bacterial plasmid DNA was extracted from a 3 ml overnight culture using the QIAprep Spin Miniprep Kit (Qiagen, 27104). Next, NGS libraries were generated following a two-step primer extension protocol.^70^ 30 μg of plasmid DNA were amplified using Kapa Hifi HotStart Ready mix (Kapa Biosystems, KK2602) in a 50 μl reaction using primers EpMap_7 and EpMap_8 (which bound to regions that were ~70 bp away from the peptide encoding region) with the following cycling parameter: 95 °C for 3 minutes, 21 cycles of 98 °C 20 seconds, 59 °C 15 seconds, 72 °C for 20 seconds and 72 °C 30 seconds final extension.

After gel purification, the final library was constructed the same way using 20 μg of PCR1 product along with primer EpMap_9 and one of 20 illumina index primers (EpMap_idx) using the following cycling parameter: 95 °C for 3 minutes, 21 cycles of 98 °C 20 seconds, 56 °C 15 seconds, 72 °C for 25 seconds and 72 °C 30 seconds final extension. Final libraries of the correct length (~400 bp) were gel-purified on a 2% (w/v) agarose-gel and subjected to fragment analyzer analysis (Advanced Analytical Technologies) using DNF-473 Standard Sensitivity NGS fragment analysis kit prior to sequencing. High-quality library pools were sequenced on the Illumina MiSeq platform using the reagent kit v3 (2×300 cycles, paired-end) with 10% PhiX.

### Bioinformatic epitope extraction

Forward and reverse reads were merged in Geneious v10.2.6 and subsequently read into R (v.4.0.4) using the ‘read.fasta’ function from the ‘seqinr’ package. The number of occurrences of each epitope within these reads was determined using the ‘str_count’ function from the ‘stringr’ package.

### Data visualization

FACS plots were created using FlowJo v10 (BD). Sequence alignments, phylogenetic trees and logo plots were exported from Geneious v10.2.6. Structural epitope visualization was performed using Pymol v2.4.2. BLI affinity traces were exported from Data Analysis HT v11.0.0.50 (FortéBio). **Fig. 1a** and **4f** were created with BioRender.com. The generation of all other figures was either already described or they were produced using Prism v8 (Graphpad).

## Author contributions

D.N., A.Y. and S.T.R. conceived the study and wrote the manuscript. D.N. and L.C. performed mouse experiments and library preparations. A.Y., A.A., R.K. performed antibody repertoire and epitope analysis. D.N. and T.C. expressed and validated antibodies. D.N., R.B.R. and T.C. performed BLI measurements. D.N. and R.A.E. performed epitope binning experiments. D.N. performed epitope mapping experiments. M.D.T. and R.A. assisted with cell sorting. S.T.R., A.O. and D.J.L. supervised the study.

## Acknowledgments

We acknowledge the ETH Zurich D-BSSE Single Cell Unit and the Genomics Facility Basel for excellent support and assistance, in particular I. Nissen, T. Schär, E. Burcklen, T. Horn and C. Beisel. We also would like to thank M.D. Hussherr and G. Camenisch for assistance with animal authorization and experiments. pB33eCPX was a gift from Patrick Daugherty (Addgene plasmid # 23336; http://n2t.net/addgene:23336; RRID:Addgene_23336). This work was supported by the European Research Council Starting Grant 679403 (to S.T.R.) and ETH Zurich Research Grants (to S.T.R.). The professorship of S.T.R. is supported by an endowment from the S. Leslie Misrock Foundation.

---

**Extended Data Figure 1.**
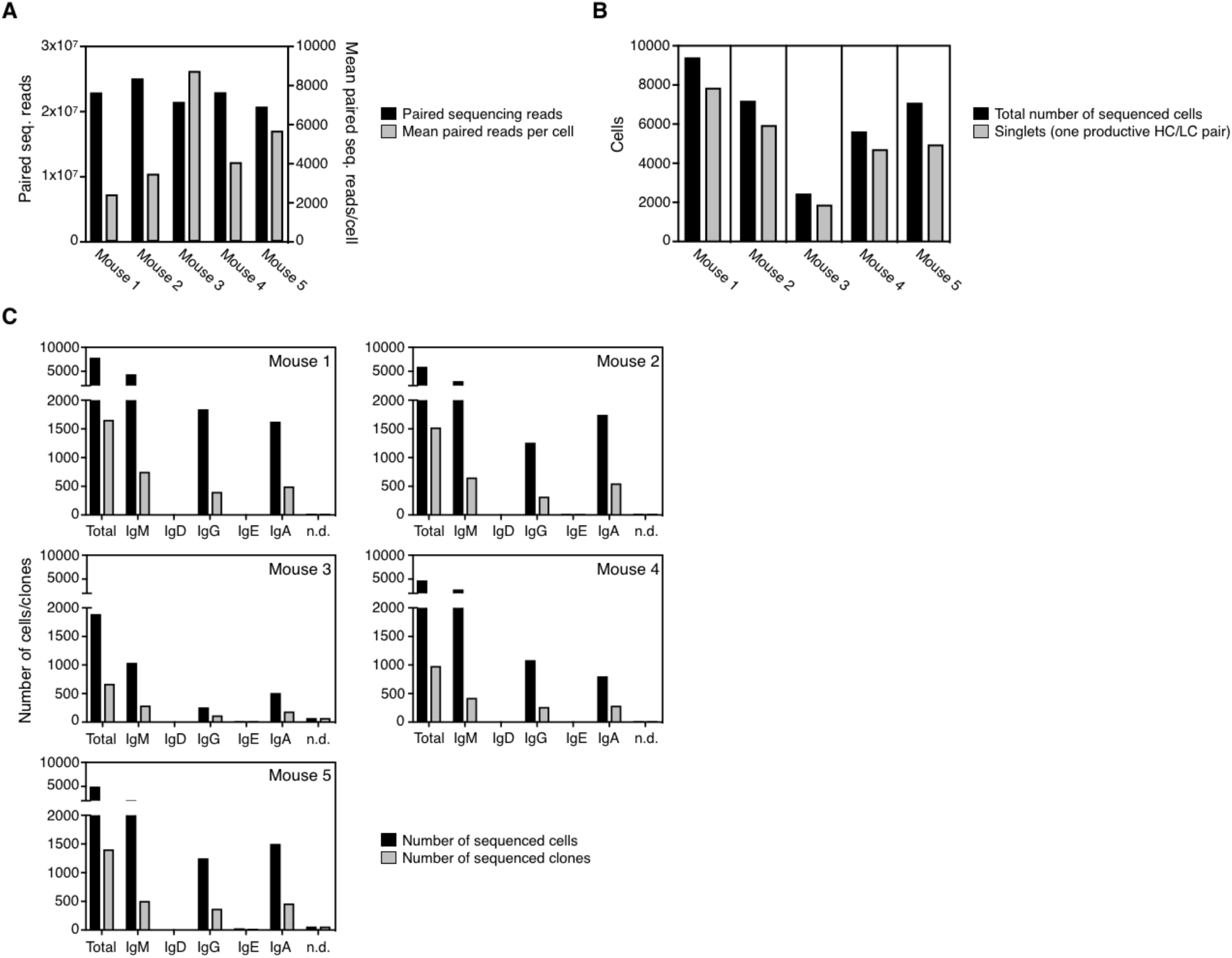
Single-cell sequencing statistics. **a.** Total number of paired sequencing reads per mouse (black, left axis) and mean paired sequencing reads per cell (grey, right axis) for all mice. **b.** Number of all sequenced cells per mouse (black) and number of cells with one productive, full-length heavy- and lightchain pair (grey). **c.** Number of sequenced productive cells (black) and clones (grey) with isotype resolution for each mouse. Clonotype definition is based on unique CDRH3-CDRL3 amino acid combination.

**Extended Data Figure 2.**
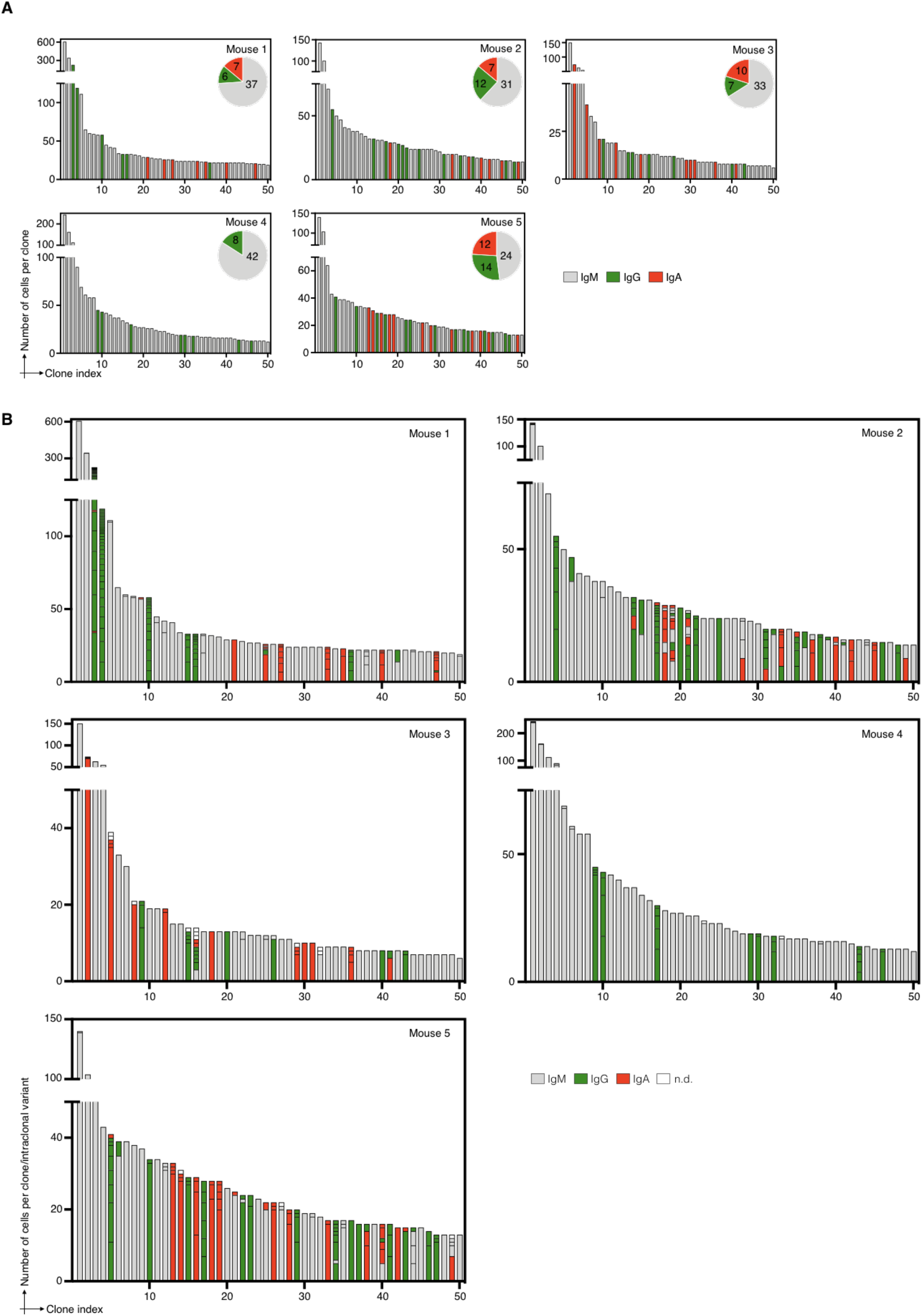
Clonal expansion profiles for the 50 most expanded clones per mouse. **a.** Clonal expansion profiles based on isotype majority. Isotypes are indicated in grey (IgM), green (IgG) and red (IgA) respectively. Pie-chart inlet indicates the numbers of isotype clones among the top 50 clones shown. **b.** Clonal expansion profiles indicating all clonal amino acid sequence variants per clone and their respective isotype assignment. Separate clonal sequence variants are shown in stacked bar plots ordered by their respective size from bottom to top. Whenever a smaller sized bar of different isotype origin intersects two bigger bars, this indicates that these cells belong to the clonal variant below. Coloring scheme according to **a** with unassigned isotype cells shown in white.

**Extended Data Figure 3.**
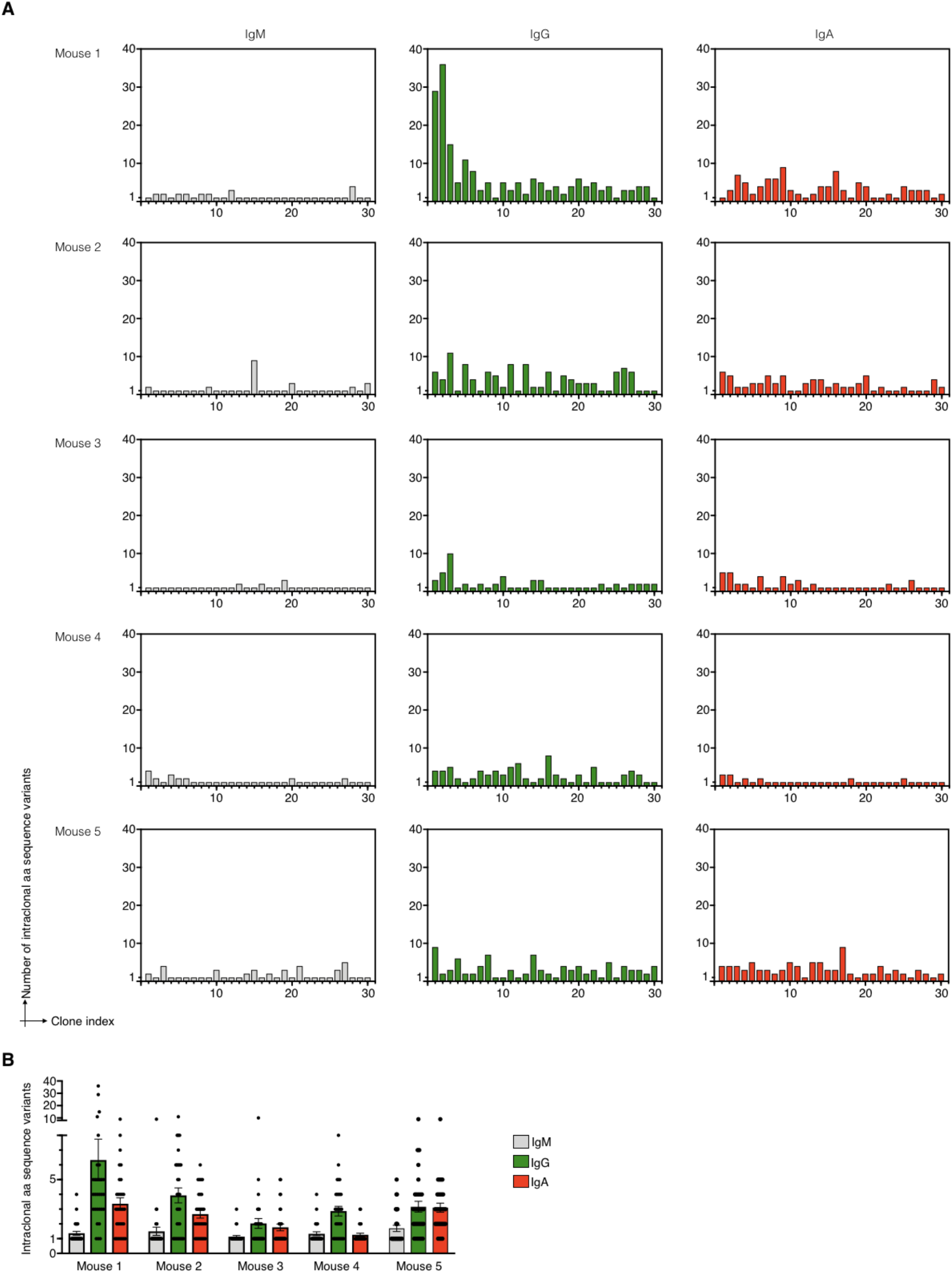
Intraclonal amino acid sequence variants for the 30 most expanded clones per isotype. **a.** Total number of intraclonal amino acid sequence variants per individual clone per isotype for all mice. **b.** Mean number of intraclonal amino acid sequence variants per isotype for all mice.

**Extended Data Figure 4.**
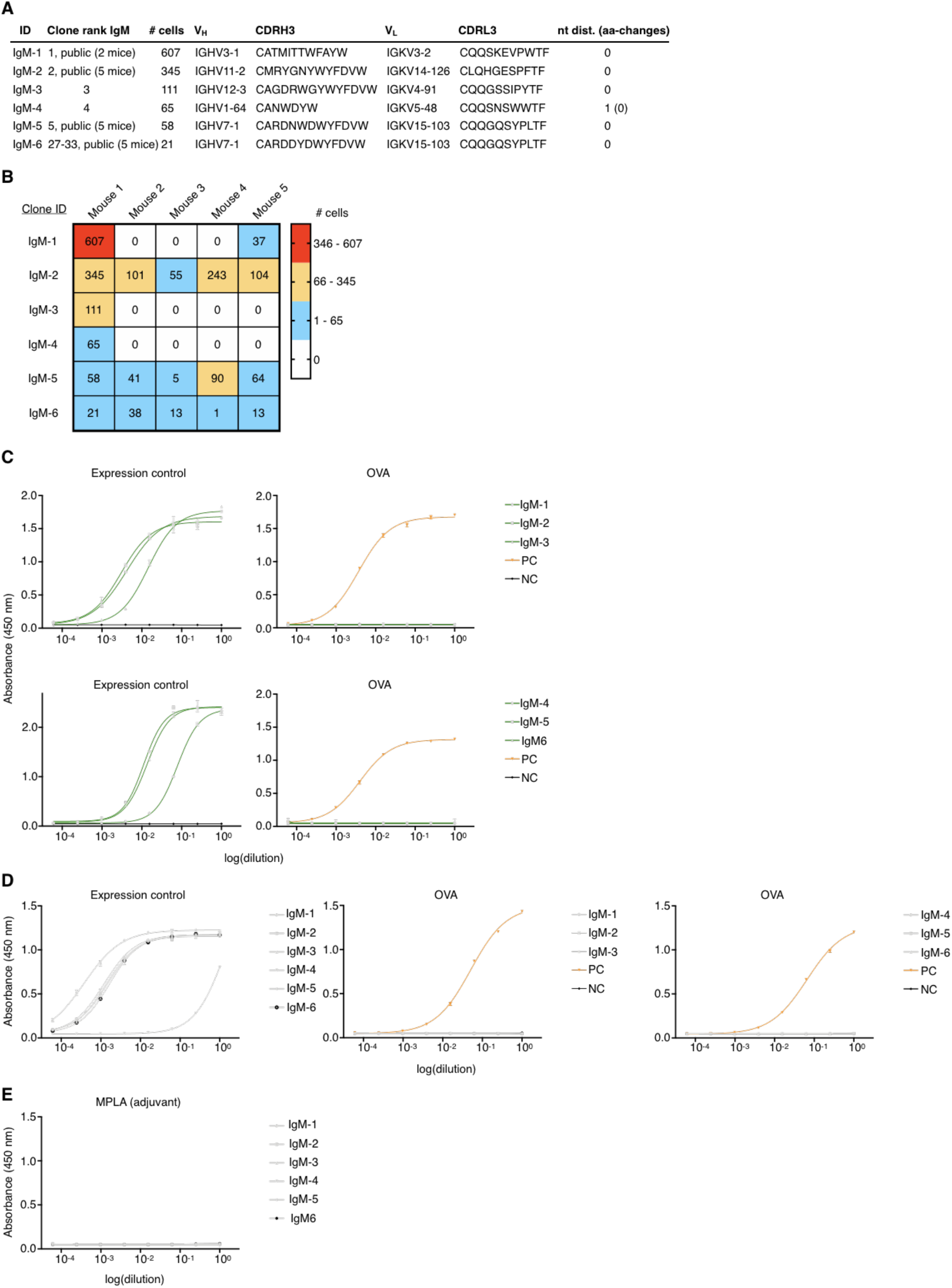
Profiling of expanded IgM clones of MS-1. **a.** Characteristics of chosen clones. **b.** Heatmap indicating the number of cells per mouse with identical clonal CDRH3-CDRL3 amino acid sequence per chosen clone. Clone ID follows **a**. **c.** Sandwich ELISA results of 3-fold serially diluted stable hybridoma cell culture supernatant of engineered cell lines expressing selected clones as IgGs. Plots on the left indicate IgG expression levels and plots on the right show binding to OVA. For each sample, two technical replicates were analysed and a four-parameter logistical curve was fitted to the data by nonlinear regression. Data are presented as the mean and error bars indicate standard deviation. Supernatant of a hybridoma cell line that does not express antibody served as negative control (NC, black) for the expression ELISA (left) and supernatant of an OVA specific inhouse cell line was used as positive control (PC, orange) for the antigen ELISA (right). **d.** Sandwich ELISA results of 3-fold serially diluted transient HEK-293 cell culture supernatant of cells expressing selected clones as IgM following **c**. Supernatant of IgM reformatted RSVF specific inhouse antibody served as negative control (NC, black) and supernatant of IgM reformatted positive clone from **c** served as positive control (PC, orange). **e.** Sandwich ELISA results on transient HEK-293 cell culture supernatant of cells expressing selected clones as IgM for binding to MPLA adjuvant.

**Extended Data Figure 5.**
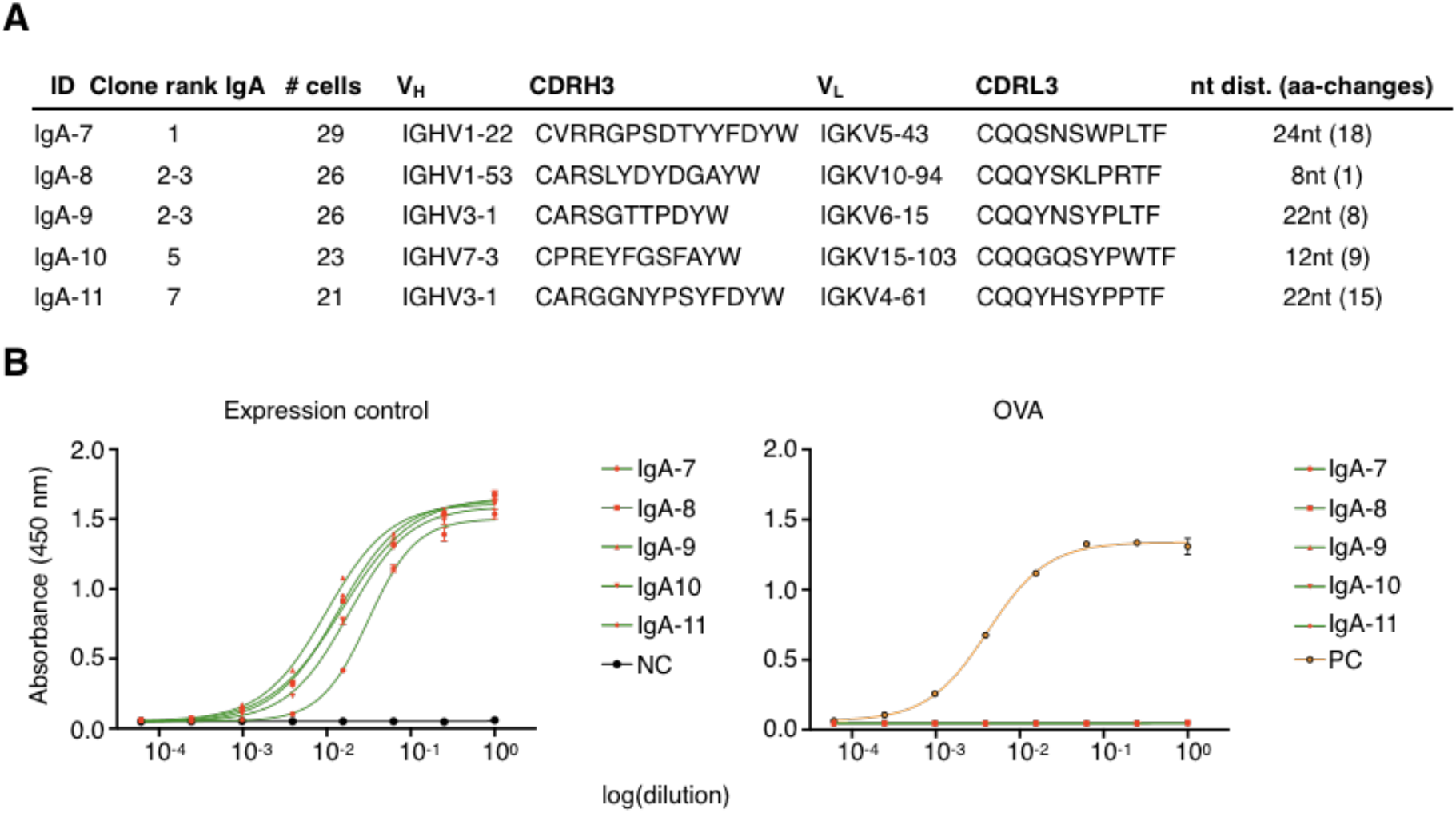
Profiling of expanded IgA clones of MS-1. **a.** Characteristics of chosen clones. **b.** Sandwich ELISA results of 3-fold serially diluted stable hybridoma cell culture supernatant of engineered cell lines expressing selected clones as IgGs. Plot on the left indicates IgG expression levels and plot on the right shows binding to OVA. For each sample, two technical replicates were analysed and a four-parameter logistical curve was fitted to the data by nonlinear regression. Data are presented as the mean and error bars indicate standard deviation. Supernatant of a hybridoma cell line that does not express antibody served as negative control (NC, black) for the expression ELISA (left) and supernatant of an OVA specific inhouse cell line was used as positive control (PC, orange) for the antigen ELISA (right).

**Extended Data Figure 6.**
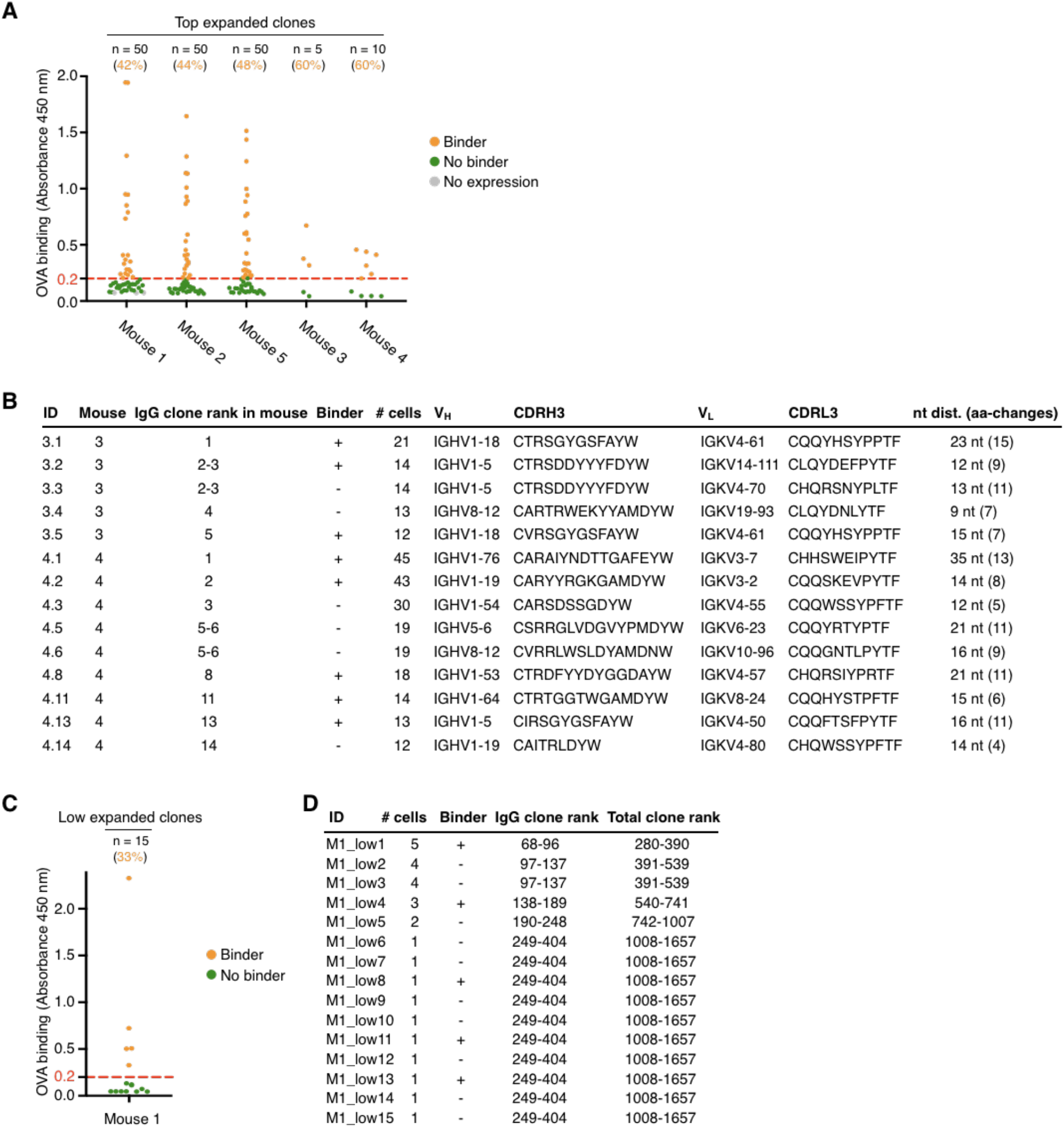
Profiling of top and low expanded IgG clones per mouse. **a.** ELISA profiling results of all top expanded clones tested per mouse. Clones were denoted binders if ELISA signal was >0.2 (3-fold above background; red dotted line). Binders are shown in orange, whereas clones that did not bind OVA or could not be expressed are shown in green and grey respectively. **b.** Overview of tested top expanded clones of MS-3 and −4. + denotes ELISA signal >0.2. **c.** ELISA profiling results of all low expanded MS-1 clones tested. **d.** Table indicating binding of low expanded clones as well as number of cells and corresponding IgG and total clone ranks within MS-1 repertoire.

**Extended Data Figure 7.**
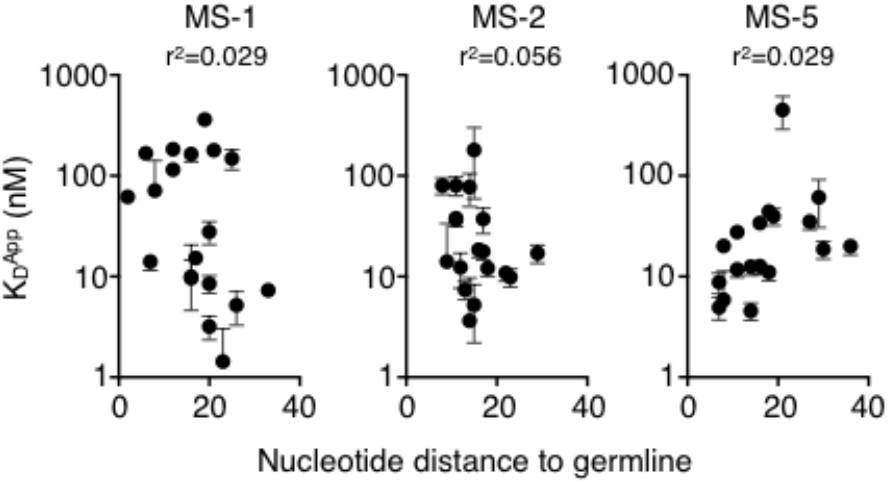
Correlation between apparent dissociation constant (K_D_^App^) and nucleotide distance to germline of clones shown in Fig. 2a, b.

**Extended Data Figure 8.**
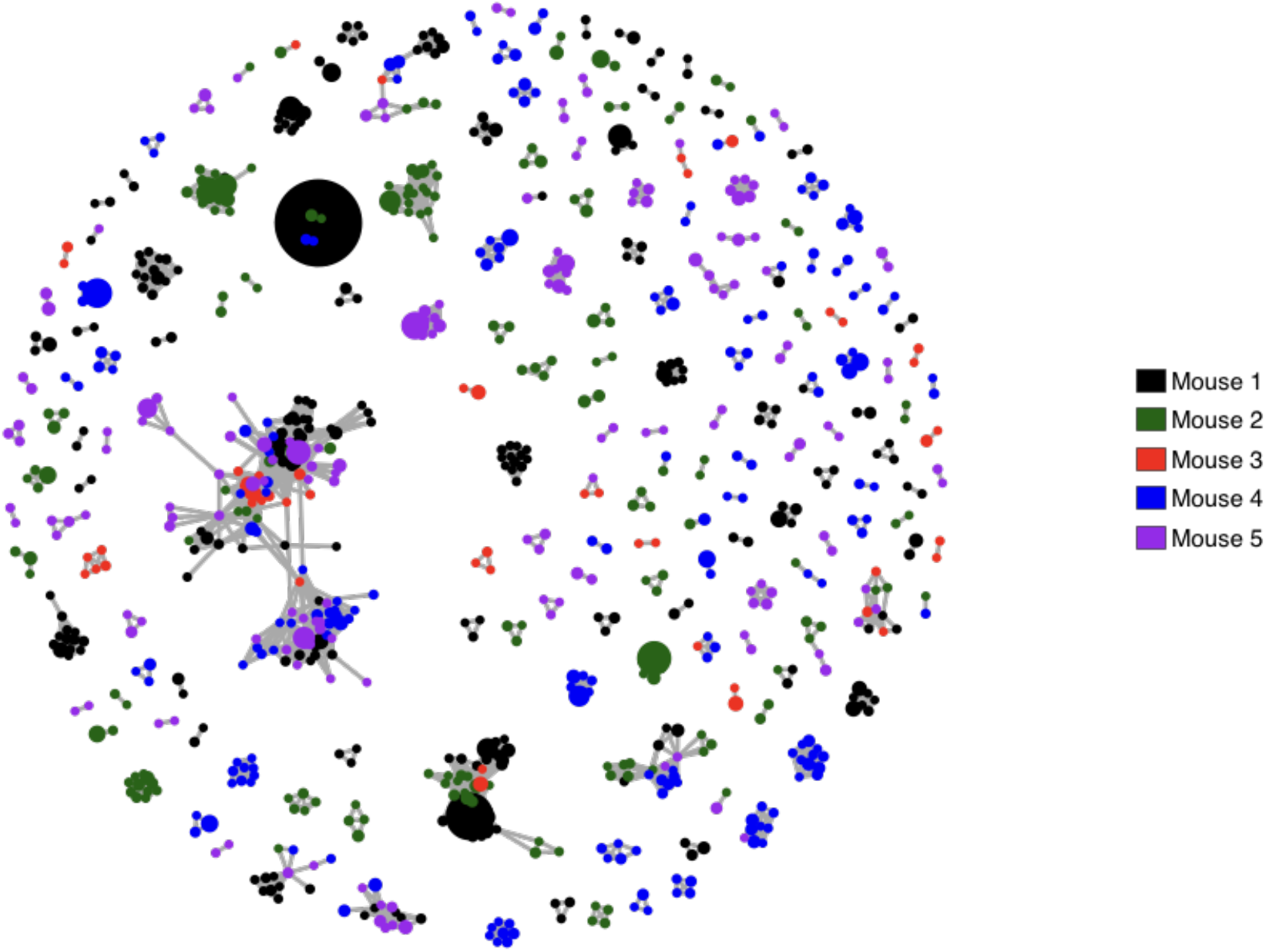
IgG similarity network. Global IgG similarity network plot for all IgG clones across all mice. Edges represent sequence nodes separated by edit distance of less than four a.a. Only those nodes with at least one edge are plotted for visualization purposes. Clones from different mice are indicated in different colors respectively. Extent of clonal expansion per clone is reflected by the size of the nodes.

**Extended Data Figure 9.**
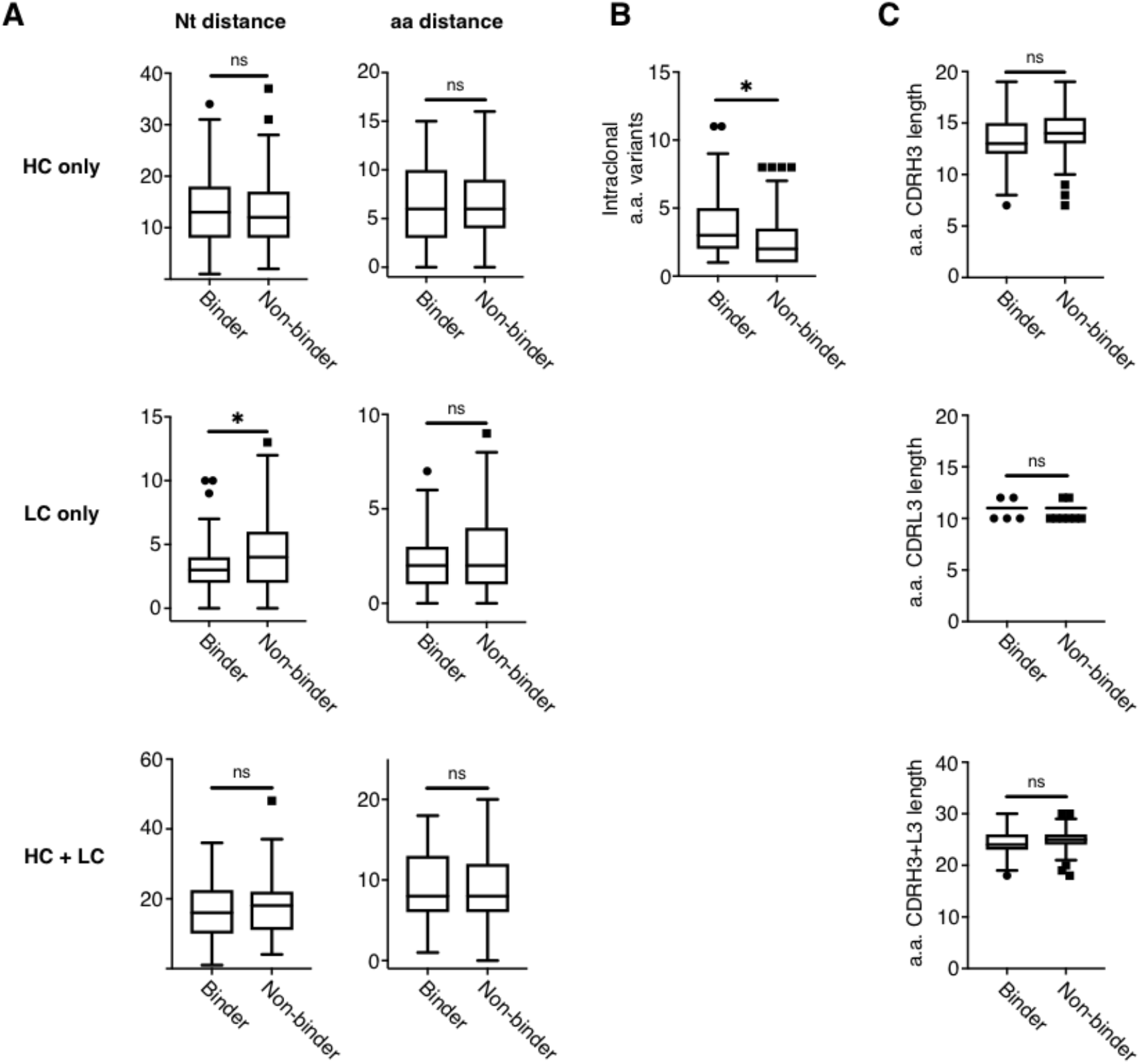
Selected metrics for binder- and non-binder pools of clones. **a.** Left: nucleotide distance to germline for heavy- and light-chain only as well as across heavy- and light-chain combined. Top: ns, not significant (P=0.70); middle: *P=0.02; bottom: ns, not significant (P=0.64); unpaired Student’s *t*-test. Right: amino acid distance to germline for heavy- and light-chain only as well as across heavy- and lightchain combined. Top: ns, not significant (P=0.77); middle: ns, not significant (P=0.11); bottom: ns, not significant (P=0.69); unpaired Student’s *t*-test. Analysis encompassed 174 experimentally verified sequences (79 binder and 95 non-binder) provided in **Extended Data Table 1** and **2** respectively. **b.** Number of intraclonal amino acid sequence variants. *P=0.02, unpaired Student’s *t*-test. Analysis contained all sequences provided in **Extended Data Table 1** and **2** except for singlet clones (76 binder and 87 non-binder). **c.** CDR3 amino acid length for CDRH3 and CDRL3 as well as CDRH3/L3 combined. Top: ns, not significant (P=0.44); middle: ns, not significant (P=0.49); bottom: ns, not significant (P=0.49); unpaired Student’s *t*-test. Analysis encompassed experimentally verified sequences (79 binder and 95 non-binder) provided in **Extended Data Table 1** and **2**.

**Extended Data Figure 10.**
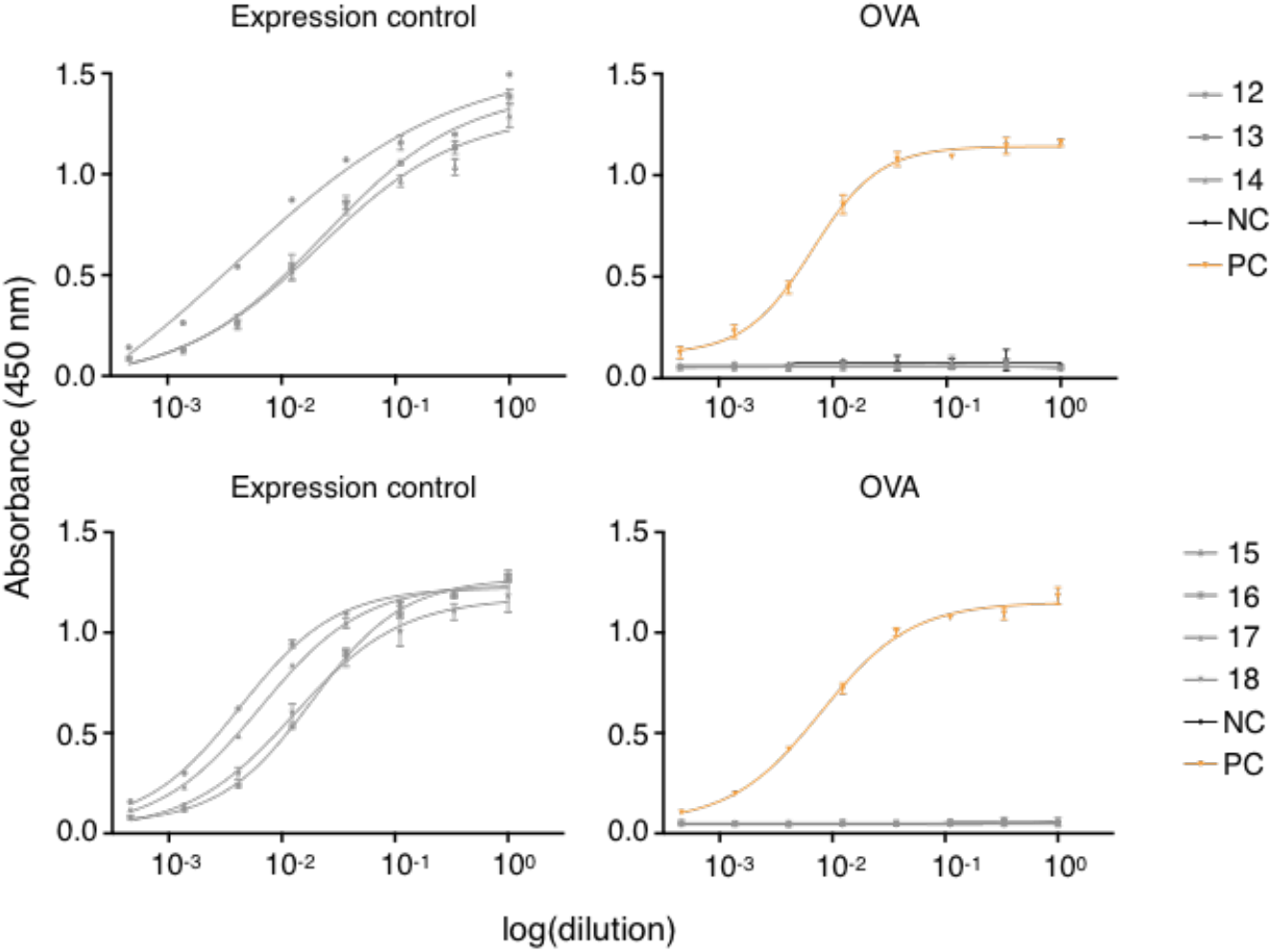
ELISA screening of multi-isotype clones shown in Fig. 2g. Sandwich ELISA results of 3-fold serially diluted stable hybridoma cell culture supernatant of engineered cell lines expressing selected clones as IgGs. Plots on the left indicate IgG expression levels and plot on the right shows binding to OVA. For each sample, two technical replicates were analysed and a four-parameter logistical curve was fitted to the data by nonlinear regression. Data are presented as the mean (n = 2 measurements) and error bars indicate standard deviation. Supernatant of a HEL specific hybridoma cell line served as negative control (NC, black) and supernatant of an OVA specific inhouse cell line was used as positive control (PC, orange).

**Extended Data Figure 11.**
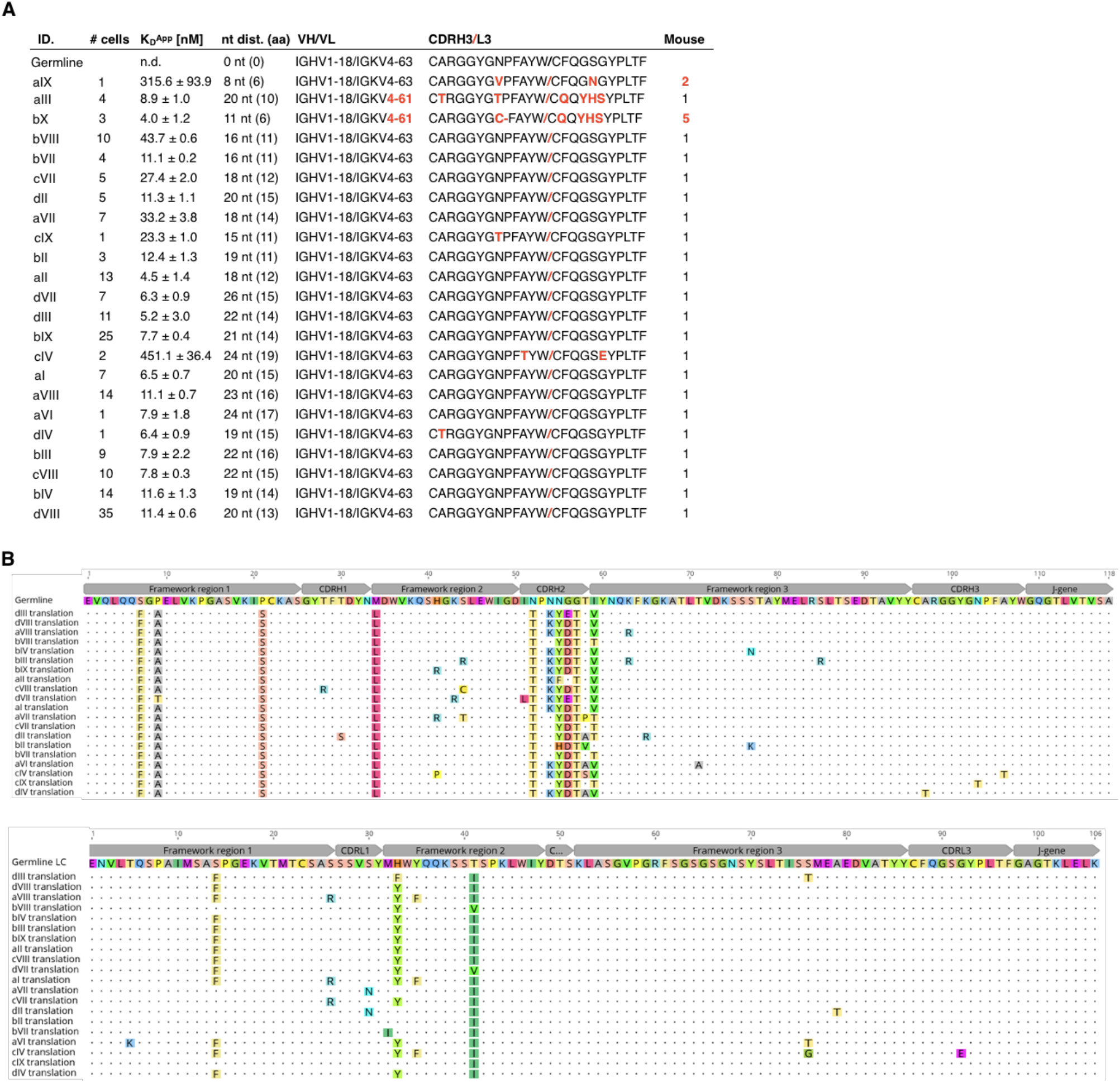
Characteristics of tested intraclonal sequence variants shown in Fig. 3b. **a.** Characteristics of tested variants. Differences in V_L_ gene usage and CDR3 amino acid sequence as well as sequences coming from different mice are indicated in red. **b.** Heavy (top) and light-chain (bottom) amino acid sequence alignment of clones with shared V/J genes. Sequence disagreements to germline are highlighted.

**Extended Data Figure 12.**
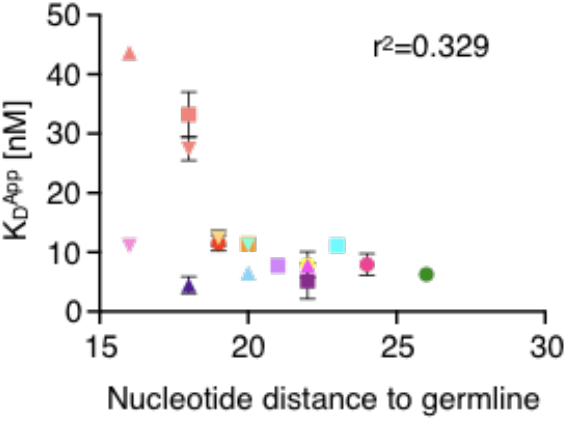
Correlation between apparent dissociation constant (K_D_^App^) and nucleotide distance to germline. Error bars indicate standard deviation (n = 3-5 measurements of K_D_).

**Extended Data Figure 13.**
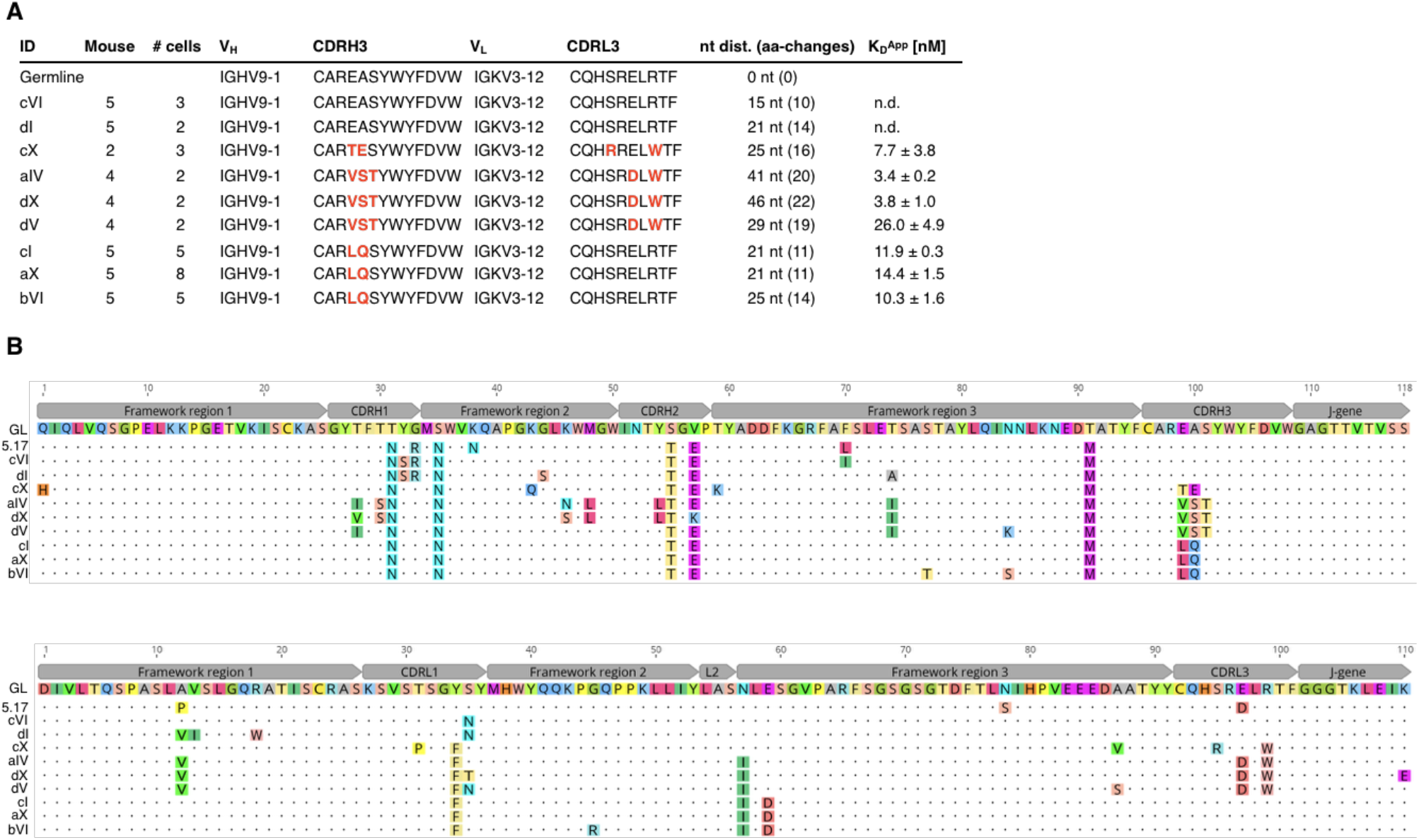
Characteristics of tested clones shown in Fig. 3e-g. **a.** Characteristics of tested clones. Differences in CDR3 amino acid sequence are indicated in red. **b.** Heavy (top) and light-chain (bottom) amino acid sequence alignment of tested clones. Sequence disagreements to germline are highlighted.

**Extended Data Figure 14.**
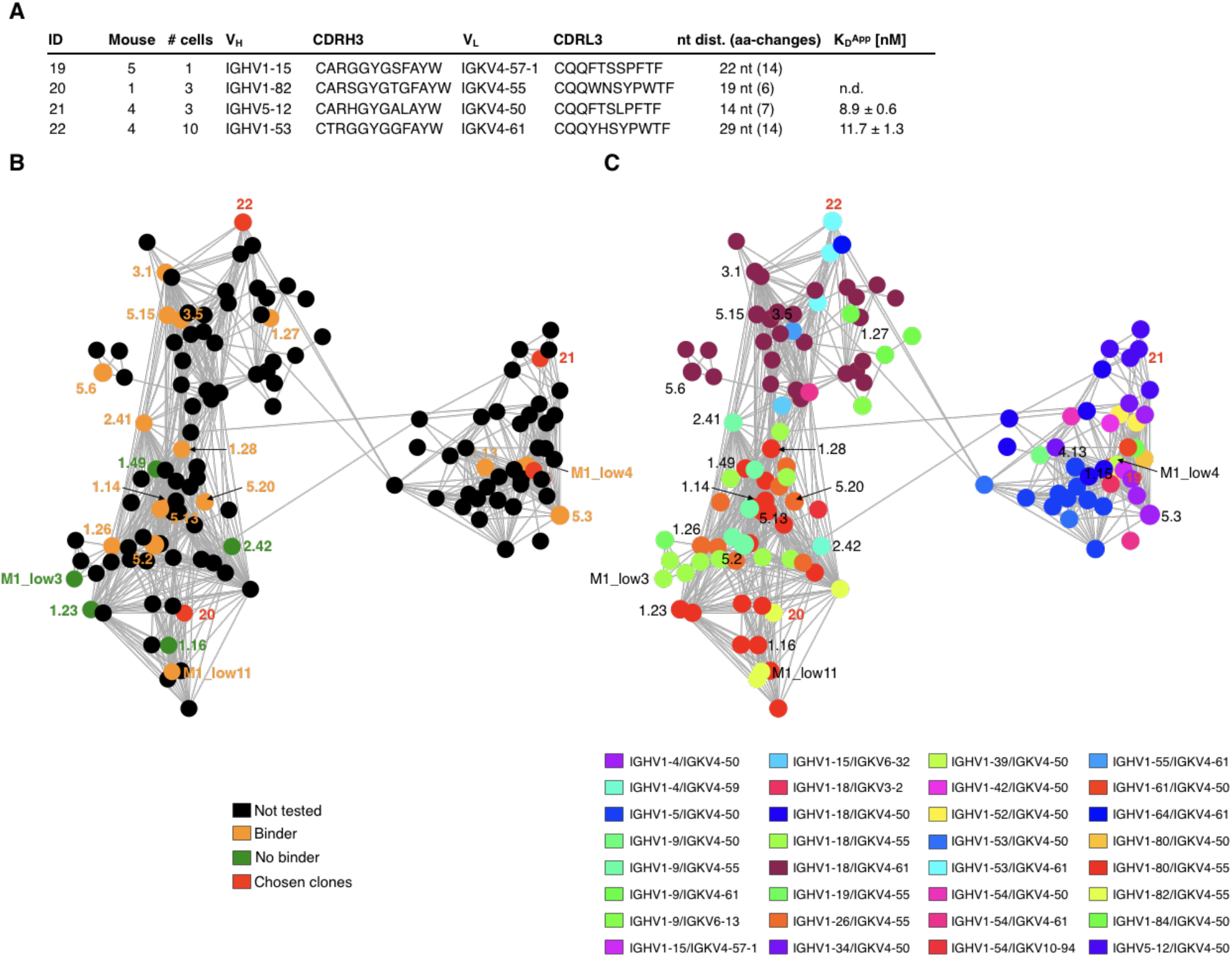
Characteristics of tested clones shown in Fig. 3h-j. **a.** Characteristics of tested clones. **b.** Subnetwork plot of connected IgG clones from all mice shown in **Extended Data Fig. 9**. Edges represent clones separated by edit distance of three or less based on the concatenated CDR3 aa sequence. Binders, non-binders, not tested clones as well as newly chosen clones are shown in orange, green, black and red respectively. Indicated clone ID according to **Extended Data Table 1** and **2**. **c.** Identical network plot as in **b**. Color code indicates differential V_H_-V_L_ gene usage as indicated.

**Extended Data Figure 15.**
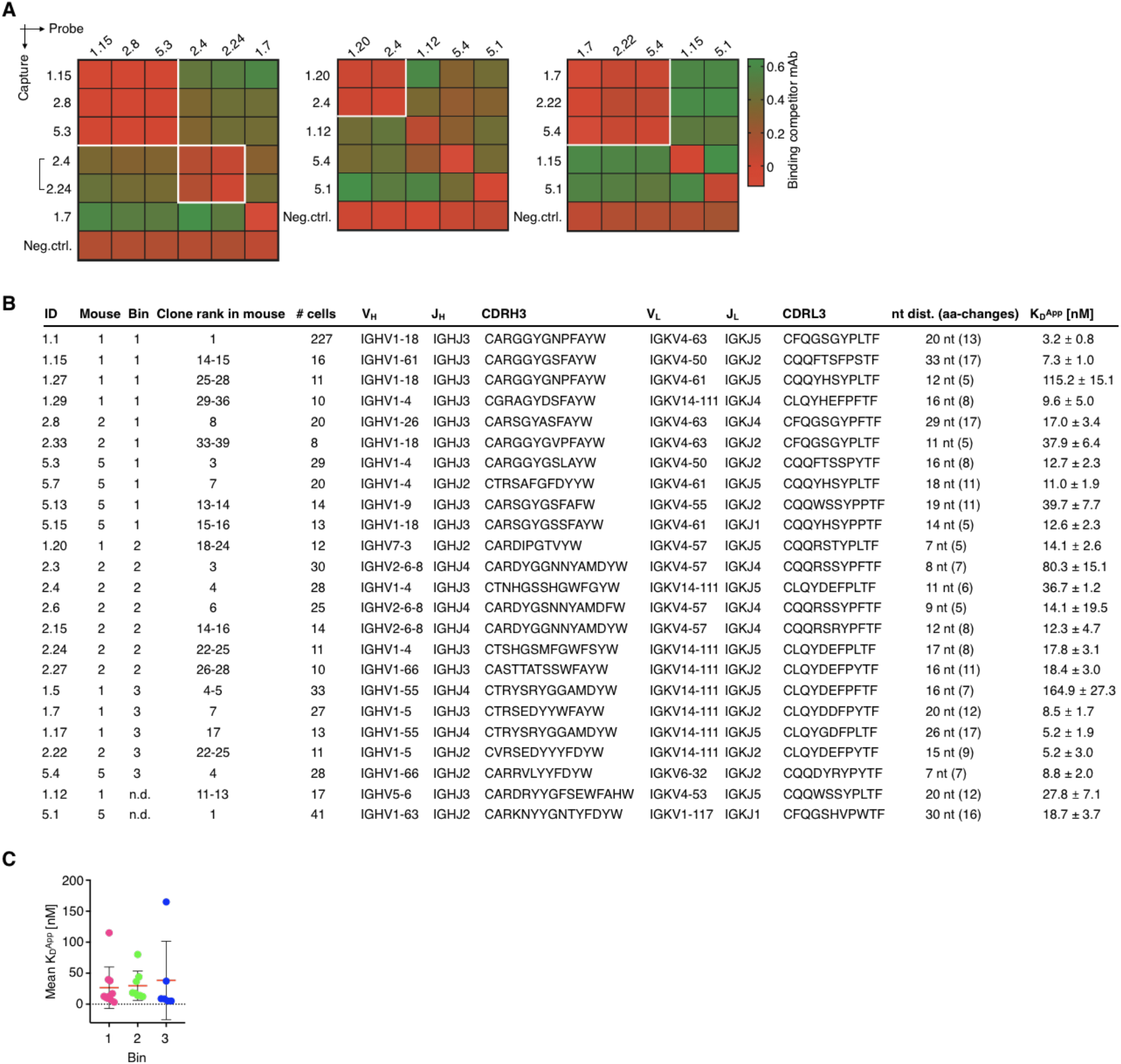
Cross-competition epitope binning between clones from different mice. **a.** Heatmaps show competitive antigen binding between top expanded, bin specific (according to **Fig. 4b**) clones from different mice. Antibodies indicated on the left were captured and probe antibodies on top were used to determine cross-competition for epitope access. Red indicates no binding of the probe antibody as a consequence of epitope blocking by the capture antibody, whereas green denotes binding of the competitor antibody. Groups of antibodies that target the same epitope (epitope bins) are highlighted in white squares. Brackets indicate clonal variants that share the same V_H_/V_L_ as well as CDR3 length and only differ in their CDR3 amino acid sequence. An anti-RSVF capture antibody, which does not bind the antigen was used as negative control for all experiments. Clone ID according to **Extended Data Table 1. b.** Characteristics of bin-specific clones. **c** Apparent dissociation constant (K_D_^App^) for binders separated by epitope bin.

**Extended Data Figure 16.**
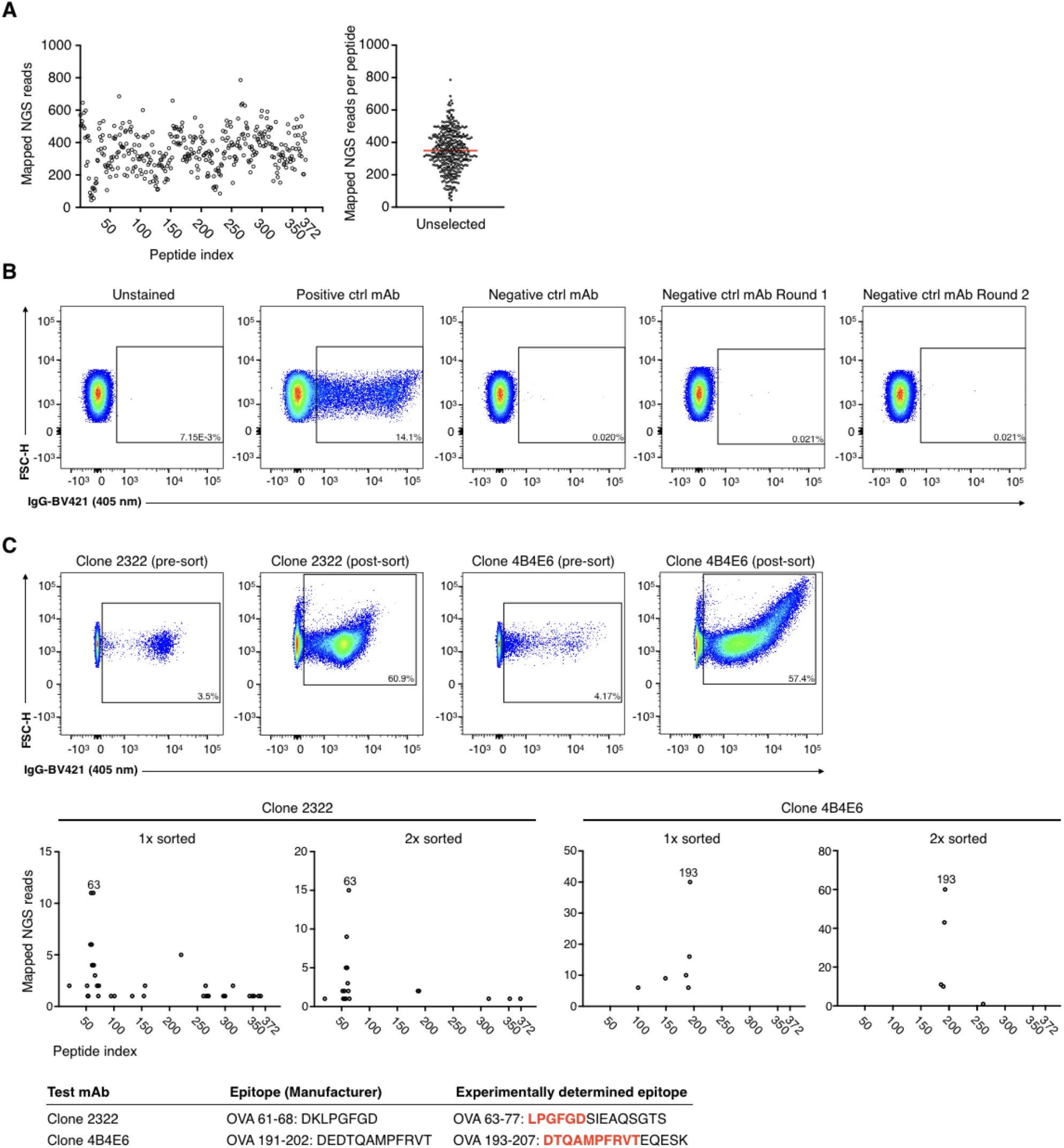
Establishment of a bacterial peptide display workflow for linear epitope mapping. **a.** Quality control of the unselected cloned OVA epitope library by NGS. All 372 peptide 15-mer windows (one amino acid offset) were observed (left) at comparable frequencies (right). Mean window occurrence for the respective NGS run is indicated in red. **b.** Flow cytometry dot plots show that secondary FACS antibody does not lead to unspecific enrichment after two rounds of enrichment. Unstained denotes no primary antibody. Positive and negative control primary mAb used were OVA specific mAb Clone 2322 (Chondrex, 7094) and an inhouse RSVF specific mouse IgG. **c.** Assay establishment with two OVA specific antibodies with known epitope specificity. Top: FACS dot plots show FACS enrichment of cells binding to commercial antibodies Clone 2322 (Chondrex, 7094) (left) and Clone 4B4E6 (Chondrex, 7096) (right). Middle: NGS results after one and two rounds of FACS enrichment. Bottom: experimentally determined epitope identity compared to epitope information provided by the manufacturer. Epitope overlap is indicated in bold red.

**Extended Data Figure 17.**
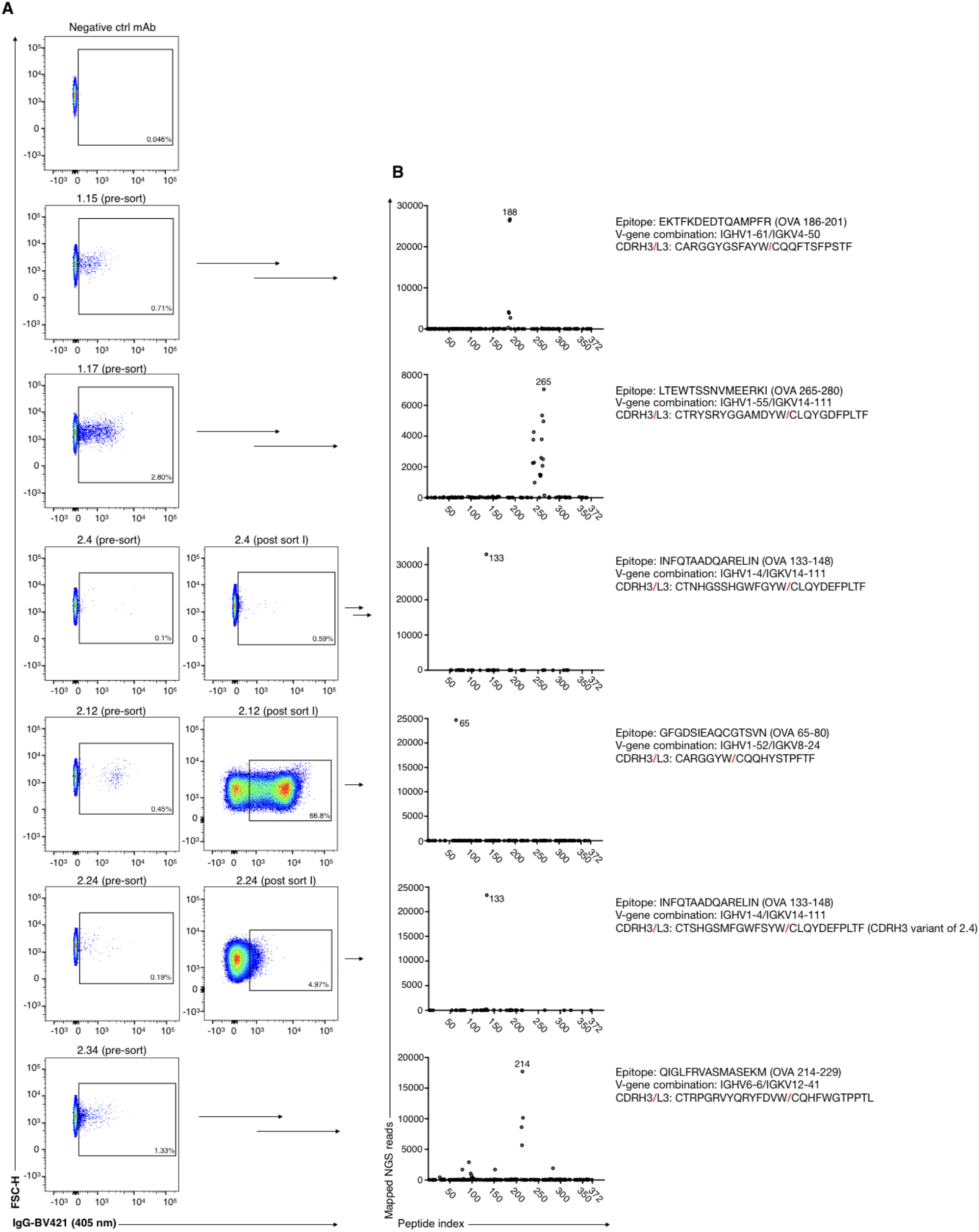
Epitope mapping results of clones with linear epitope specificity as shown in Fig. 4e. **a.** Flow cytometry dot plots indicate FACS enrichment of positive clones for each antibody. **b.** Deep sequencing results of positive FACS output and epitope assignment for each antibody. Numbers correspond to the peptide index that was enriched the most. Only peptides with one or more occurrences are shown.

**Extended Data Table 1.**
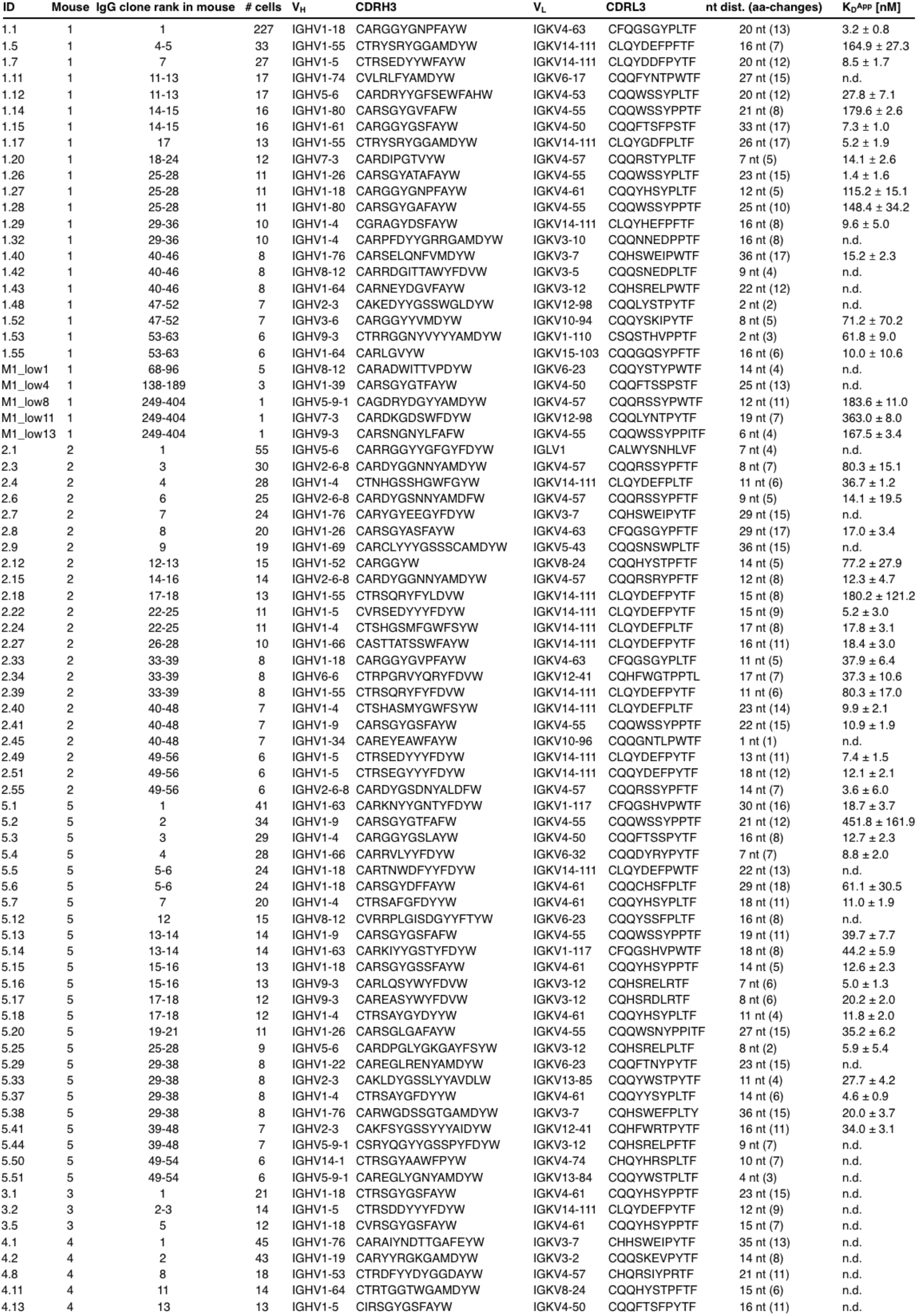
Overview and characteristics of OVA binders. Clone ID corresponds to mouse number followed by clone index number corresponding to **Fig. 1g**.

| ID | Mouse | IgG clone rank in mouse | # cells | V<sub>H</sub> | CDRH3 | V<sub>L</sub> | CDRL3 | nt dist. (aa-changes) | K<sub>D</sub><sup>APP</sup> [nM] |
| --- | --- | --- | --- | --- | --- | --- | --- | --- | --- |
| 1.1 | 1 | 1 | 227 | IGHV1-18 | CARGGGYNPFAYW | IGKV4-63 | CFQGGSGYPLTF | 20 nt (13) | 3.2 ± 0.8 |
| 1.5 | 1 | 4-5 | 33 | IGHV1-55 | CTRYSTRYGGAMDYW | IGKV14-111 | CLQYDEFFPFTF | 16 nt (7) | 164.9 ± 27.3 |
| 1.7 | 1 | 7 | 27 | IGHV1-5 | CTRSEDIYWFAYW | IGKV14-111 | CLQYDDFPYTF | 20 nt (12) | 8.5 ± 1.7 |
| 1.11 | 1 | 11-13 | 17 | IGHV1-74 | CVLRIFYAMDYW | IGKV6-17 | CQQFYNTPWTF | 27 nt (15) | n.d. |
| 1.12 | 1 | 11-13 | 17 | IGHV5-6 | CARDRYYGSEWFAHW | IGKV4-53 | CQQWSSYPPLTF | 20 nt (12) | 27.8 ± 7.1 |
| 1.14 | 1 | 14-15 | 16 | IGHV1-80 | CARSGYGVFAFW | IGKV4-55 | CQQWSSYPPTF | 21 nt (8) | 179.6 ± 2.6 |
| 1.15 | 1 | 14-15 | 16 | IGHV1-61 | CARGGYGSFAYW | IGKV4-50 | CQQFTSFPSTF | 33 nt (17) | 7.3 ± 1.0 |
| 1.17 | 1 | 17 | 13 | IGHV1-55 | CTRYSTRYGGAMDYW | IGKV14-111 | CLQYGDFFPLTF | 26 nt (17) | 5.2 ± 1.9 |
| 1.20 | 1 | 18-24 | 12 | IGHV7-3 | CARDIPGTVYW | IGKV4-57 | CQQRSTYPLTF | 7 nt (5) | 14.1 ± 2.6 |
| 1.26 | 1 | 25-28 | 11 | IGHV1-26 | CARSGYATFAYW | IGKV4-55 | CQQWSSYPPLTF | 23 nt (15) | 1.4 ± 1.6 |
| 1.27 | 1 | 25-28 | 11 | IGHV1-18 | CARGGYGNPFAYW | IGKV4-61 | CQQYHSYPLTF | 12 nt (5) | 115.2 ± 15.1 |
| 1.28 | 1 | 25-28 | 11 | IGHV1-80 | CARSGYGAFAYW | IGKV4-55 | CQQWSSYPPTF | 25 nt (10) | 148.4 ± 34.2 |
| 1.29 | 1 | 29-36 | 10 | IGHV1-4 | CGRAGYDSFAYW | IGKV14-111 | CLQYHEFFPFTF | 16 nt (8) | 9.6 ± 5.0 |
| 1.32 | 1 | 29-36 | 10 | IGHV1-4 | CARPFDYGGRRGAMDYW | IGKV3-10 | CQQNNEDPPTF | 16 nt (8) | n.d. |
| 1.40 | 1 | 40-46 | 8 | IGHV1-76 | CARSELQNFVMDYW | IGKV3-7 | CQHSWEIPWTF | 36 nt (17) | 15.2 ± 2.3 |
| 1.42 | 1 | 40-46 | 8 | IGHV8-12 | CARRDGITTAWFYDWW | IGKV3-5 | CQQSNEDPLTF | 9 nt (4) | n.d. |
| 1.43 | 1 | 40-46 | 8 | IGHV1-64 | CARNEYDGVFAYW | IGKV3-12 | CQHSRELPTWTF | 22 nt (12) | n.d. |
| 1.48 | 1 | 47-52 | 7 | IGHV2-3 | CAKEDYYGSSWGLDYW | IGKV12-98 | CQQLYSTPYTF | 2 nt (2) | n.d. |
| 1.52 | 1 | 47-52 | 7 | IGHV3-6 | CARGGYVMDYW | IGKV10-94 | CQQYSKIPYTF | 8 nt (5) | 71.2 ± 70.2 |
| 1.53 | 1 | 53-63 | 6 | IGHV9-3 | CTRRGGNYVYYYAMDYW | IGKV1-110 | CSQSTHVPPTF | 2 nt (3) | 61.8 ± 9.0 |
| 1.55 | 1 | 53-63 | 6 | IGHV1-64 | CARLGYYW | IGKV15-103 | CQQGQSPPTF | 16 nt (6) | 10.0 ± 10.6 |
| M1_low1 | 1 | 68-96 | 5 | IGHV8-12 | CARADWITTPDYW | IGKV6-23 | CQQYSTYPWTF | 14 nt (4) | n.d. |
| M1_low4 | 1 | 138-189 | 3 | IGHV1-39 | CARSGYGTFAFW | IGKV4-50 | CQQFTSSPSTF | 25 nt (13) | n.d. |
| M1_low8 | 1 | 249-404 | 1 | IGHV5-9-1 | CAGDRYDGYAMDYW | IGKV4-57 | CQQRSSYPWTF | 12 nt (11) | 183.6 ± 11.0 |
| M1_low11 | 1 | 249-404 | 1 | IGHV7-3 | CARDKGDWFDYW | IGKV12-98 | CQQLYNTPYTF | 19 nt (7) | 363.0 ± 8.0 |
| M1_low13 | 1 | 249-404 | 1 | IGHV9-3 | CARSNGNYLFAFW | IGKV4-55 | CQQWSSYPPTF | 6 nt (4) | 167.5 ± 3.4 |
| 2.1 | 2 | 1 | 55 | IGHV5-6 | CARRGGYGFYDYYW | IGLV1 | CALWYNNHLVF | 7 nt (4) | n.d. |
| 2.3 | 2 | 3 | 30 | IGHV2-6-8 | CARDYGGNNYAMDYW | IGKV4-57 | CQQRSSYPPTF | 8 nt (7) | 80.3 ± 15.1 |
| 2.4 | 2 | 4 | 28 | IGHV1-4 | CTNHGSSHGWFYW | IGKV14-111 | CLQYDEFFPLTF | 11 nt (6) | 36.7 ± 1.2 |
| 2.6 | 2 | 6 | 25 | IGHV2-6-8 | CARDYGSNNYAMDFW | IGKV4-57 | CQQRSSYPPTF | 9 nt (5) | 14.1 ± 19.5 |
| 2.7 | 2 | 7 | 24 | IGHV1-76 | CARYGYEEGYFDYW | IGKV3-7 | CQHSWEIPYTF | 29 nt (15) | n.d. |
| 2.8 | 2 | 8 | 20 | IGHV1-26 | CARSGYASFAYW | IGKV4-63 | CFQGGSGYPPTF | 29 nt (17) | 17.0 ± 3.4 |
| 2.9 | 2 | 9 | 19 | IGHV1-69 | CARCLYYGSSSCAMDYW | IGKV5-43 | CQQSNSWPLTF | 36 nt (15) | n.d. |
| 2.12 | 2 | 12-13 | 15 | IGHV1-52 | CARGGYW | IGKV8-24 | CQQHYSTPPTF | 14 nt (5) | 77.2 ± 27.9 |
| 2.15 | 2 | 14-16 | 14 | IGHV2-6-8 | CARDYGGNNYAMDYW | IGKV4-57 | CQQRSSYPPTF | 12 nt (8) | 12.3 ± 4.7 |
| 2.18 | 2 | 17-18 | 13 | IGHV1-55 | CTRSQRYFYLDVW | IGKV14-111 | CLQYDEFFPYTF | 15 nt (8) | 180.2 ± 121.2 |
| 2.22 | 2 | 22-25 | 11 | IGHV1-5 | CVRSEDIYFYDYYW | IGKV14-111 | CLQYDEFFPYTF | 15 nt (9) | 5.2 ± 3.0 |
| 2.24 | 2 | 22-25 | 11 | IGHV1-4 | CTSHGSMFGWFSYW | IGKV14-111 | CLQYDEFFPLTF | 17 nt (8) | 17.8 ± 3.1 |
| 2.27 | 2 | 26-28 | 10 | IGHV1-66 | CATTATSSWFAYW | IGKV14-111 | CLQYDEFFPYTF | 16 nt (11) | 18.4 ± 3.0 |
| 2.33 | 2 | 33-39 | 8 | IGHV1-18 | CARGGYGVPFAYW | IGKV4-63 | CFQGGSGYPLTF | 11 nt (5) | 37.9 ± 6.4 |
| 2.34 | 2 | 33-39 | 8 | IGHV6-6 | CTRPGRVYQRYFDVW | IGKV12-41 | CQHFWGTPTPL | 17 nt (7) | 37.3 ± 10.6 |
| 2.39 | 2 | 33-39 | 8 | IGHV1-55 | CTRSQRYFYFDVW | IGKV14-111 | CLQYDEFFPYTF | 11 nt (6) | 80.3 ± 17.0 |
| 2.40 | 2 | 40-48 | 7 | IGHV1-4 | CTSHASMYGWFSYW | IGKV14-111 | CLQYDEFFPLTF | 23 nt (14) | 9.9 ± 2.1 |
| 2.41 | 2 | 40-48 | 7 | IGHV1-9 | CARSGYGSFAYW | IGKV4-55 | CQQWSSYPPTF | 22 nt (15) | 10.9 ± 1.9 |
| 2.45 | 2 | 40-48 | 7 | IGHV1-34 | CAREYEAFFAYW | IGKV10-96 | CQQGNTLPWTF | 1 nt (1) | n.d. |
| 2.49 | 2 | 49-56 | 6 | IGHV1-5 | CTRSEDIYFYDYYW | IGKV14-111 | CLQYDEFFPYTF | 13 nt (11) | 7.4 ± 1.5 |
| 2.51 | 2 | 49-56 | 6 | IGHV1-5 | CTRSEGYFYDYYW | IGKV14-111 | CQQYDEFFPYTF | 18 nt (12) | 12.1 ± 2.1 |
| 2.55 | 2 | 49-56 | 6 | IGHV2-6-8 | CARDYGSNDYALDFW | IGKV4-57 | CQQRSSYPPTF | 14 nt (7) | 3.6 ± 6.0 |
| 5.1 | 5 | 1 | 41 | IGHV1-63 | CARKIYYGNTYFDYW | IGKV1-117 | CFQGSHVPWTF | 30 nt (16) | 18.7 ± 3.7 |
| 5.2 | 5 | 2 | 34 | IGHV1-9 | CARSGYGTFAFW | IGKV4-55 | CQQWSSYPPTF | 21 nt (12) | 451.8 ± 161.9 |
| 5.3 | 5 | 3 | 29 | IGHV1-4 | CARGGYGSLAYW | IGKV4-50 | CQQFTSSPYTF | 16 nt (8) | 12.7 ± 2.3 |
| 5.4 | 5 | 4 | 28 | IGHV1-66 | CARRVLYFYDYYW | IGKV6-32 | CQQDYRYPYTF | 7 nt (7) | 8.8 ± 2.0 |
| 5.5 | 5 | 5-6 | 24 | IGHV1-18 | CARTNWDFYFYDYYW | IGKV14-111 | CLQYDEFFWTF | 22 nt (13) | n.d. |
| 5.6 | 5 | 5-6 | 24 | IGHV1-18 | CARSGYDFFAYW | IGKV4-61 | CQQCHSFPLTF | 29 nt (18) | 61.1 ± 30.5 |
| 5.7 | 5 | 7 | 20 | IGHV1-4 | CTRSAGFDYDYYW | IGKV4-61 | CQQYHSYPLTF | 18 nt (11) | 11.0 ± 1.9 |
| 5.12 | 5 | 12 | 15 | IGHV8-12 | CVRRLGISDGYFYTYW | IGKV6-23 | CQQYSSFPLTF | 16 nt (8) | n.d. |
| 5.13 | 5 | 13-14 | 14 | IGHV1-9 | CARSGYGSFAFW | IGKV4-55 | CQQWSSYPPTF | 19 nt (11) | 39.7 ± 7.7 |
| 5.14 | 5 | 13-14 | 14 | IGHV1-63 | CARKIYYGSTYFDYW | IGKV1-117 | CFQGSHVPWTF | 18 nt (8) | 44.2 ± 5.9 |
| 5.15 | 5 | 15-16 | 13 | IGHV1-18 | CARSGYGSFAYW | IGKV4-61 | CQQYHSYPPTF | 14 nt (5) | 12.6 ± 2.3 |
| 5.16 | 5 | 15-16 | 13 | IGHV9-3 | CARLQSYWYFDVW | IGKV3-12 | CQHSRELRTF | 7 nt (6) | 5.0 ± 1.3 |
| 5.17 | 5 | 17-18 | 12 | IGHV9-3 | CAREASYWYFDVW | IGKV3-12 | CQHSRDLRTF | 8 nt (6) | 20.2 ± 2.0 |
| 5.18 | 5 | 17-18 | 12 | IGHV1-4 | CTRSAYGYDYYW | IGKV4-61 | CQQYHSYPLTF | 11 nt (4) | 11.8 ± 2.0 |
| 5.20 | 5 | 19-21 | 11 | IGHV1-26 | CARSLGAFAYW | IGKV4-55 | CQQWSNYPPTF | 27 nt (15) | 35.2 ± 6.2 |
| 5.25 | 5 | 25-28 | 9 | IGHV5-6 | CARDPGLYGKGYFSYW | IGKV3-12 | CQHSRELPLTF | 8 nt (2) | 5.9 ± 5.4 |
| 5.29 | 5 | 29-38 | 8 | IGHV1-22 | CAREGLRENYAMDYW | IGKV6-23 | CQQFTNYPYTF | 23 nt (15) | n.d. |
| 5.33 | 5 | 29-38 | 8 | IGHV2-3 | CAKLDYGSLLYAVDLW | IGKV13-85 | CQQYWSYPYTF | 11 nt (4) | 27.7 ± 4.2 |
| 5.37 | 5 | 29-38 | 8 | IGHV1-4 | CTRSAYGFDYDYYW | IGKV4-61 | CQQYHSYPLTF | 14 nt (6) | 4.6 ± 0.9 |
| 5.38 | 5 | 29-38 | 8 | IGHV1-76 | CARWGDSSGTGAMDYW | IGKV3-7 | CQHSWEFPLTY | 36 nt (15) | 20.0 ± 3.7 |
| 5.41 | 5 | 39-48 | 7 | IGHV2-3 | CAKFSYGSYYAIDYW | IGKV12-41 | CQHFWRTPYTF | 16 nt (11) | 34.0 ± 3.1 |
| 5.44 | 5 | 39-48 | 7 | IGHV5-9-1 | CSRYQGGYGSPPYFDYW | IGKV3-12 | CQHSRELPTF | 9 nt (7) | n.d. |
| 5.50 | 5 | 49-54 | 6 | IGHV14-1 | CTRSYAAWFPYW | IGKV4-74 | CHQYHRSPLTF | 10 nt (7) | n.d. |
| 5.51 | 5 | 49-54 | 6 | IGHV5-9-1 | CAREGLYGNAMDYW | IGKV13-84 | CQQYWSYPLTF | 4 nt (3) | n.d. |
| 3.1 | 3 | 1 | 21 | IGHV1-18 | CTRSYGSFAYW | IGKV4-61 | CQQYHSYPPTF | 23 nt (15) | n.d. |
| 3.2 | 3 | 2-3 | 14 | IGHV1-5 | CTRSDDYFYDYYW | IGKV14-111 | CLQYDEFFPYTF | 12 nt (9) | n.d. |
| 3.5 | 3 | 5 | 12 | IGHV1-18 | CVRSGYGSFAYW | IGKV4-61 | CQQYHSYPPTF | 15 nt (7) | n.d. |
| 4.1 | 4 | 1 | 45 | IGHV1-76 | CARAIYNDTTGAFYEW | IGKV3-7 | CHHSWEIPYTF | 35 nt (13) | n.d. |
| 4.2 | 4 | 2 | 43 | IGHV1-19 | CARYRGKGAMDYW | IGKV3-2 | CQKSKEVPYTF | 14 nt (8) | n.d. |
| 4.8 | 4 | 8 | 18 | IGHV1-53 | CTRDFYDYGGDAYW | IGKV4-57 | CHQRSIYPTF | 21 nt (11) | n.d. |
| 4.11 | 4 | 11 | 14 | IGHV1-64 | CTRTGGTWGAMDYW | IGKV8-24 | CQQHYSTPPTF | 15 nt (6) | n.d. |
| 4.13 | 4 | 13 | 13 | IGHV1-5 | CIRSGYGSFAYW | IGKV4-50 | CQQFTSFPYTF | 16 nt (11) | n.d. |

**Extended Data Table 2.** Overview and characteristics of non-binders. Clone ID corresponds to mouse number followed by clone index number corresponding to **Fig. 1g**.

| ID | Mouse | IgG clone rank in mouse | # cells | V<sub>H</sub> | CDRH3 | V<sub>L</sub> | CDRL3 | nt dist. (aa-changes) |
| --- | --- | --- | --- | --- | --- | --- | --- | --- |
| 1.6 | 1 | 6 | 31 | IGHV5-9-1 | CTGDRYDGYAMDYW | IGKV4-57 | CQQRSSYPWTF | 6 nt (4) |
| 1.8 | 1 | 8 | 22 | IGHV10-1 | CVRQGGKIYYDYDFDYW | IGKV17-127 | CLQSDNMPYTF | 4 nt (2) |
| 1.9 | 1 | 9-10 | 18 | IGHV9-3 | CTRRGGNYYVYAMDYW | IGKV14-111 | CLQYDELYTF | 4 nt (4) |
| 1.10 | 1 | 9-10 | 18 | IGHV9-3 | CARSDGNLYFAYW | IGKV4-55 | CQQWSSYPPTF | 4 nt (1) |
| 1.13 | 1 | 11-13 | 17 | IGHV1-34 | CARSLYDGYPHFDYW | IGKV4-50 | CQQTSSPPTF | 11 nt (8) |
| 1.16 | 1 | 16 | 14 | IGHV1-80 | CARSGYGTGFAYW | IGKV4-55 | CQQWNSYPWTF | 19 nt (6) |
| 1.18 | 1 | 18-24 | 12 | IGHV1-5 | CTRSDSNFYFDYW | IGKV14-111 | CLQYDEFPTYF | 11 nt (7) |
| 1.19 | 1 | 18-24 | 12 | IGHV5-6 | CARYDNYVRVAMDYW | IGKV4-74 | CHQYHRSPTYF | 18 nt (7) |
| 1.21 | 1 | 18-24 | 12 | IGHV13-2 | CSRAFHYYDYGYMDYW | IGKV9-124 | CLQYASYPWTF | 13 nt (9) |
| 1.23 | 1 | 18-24 | 12 | IGHV1-80 | CARSGYGTGFAYW | IGKV4-55 | CQQWSSYPWTF | 18 nt (6) |
| 1.25 | 1 | 25-28 | 11 | IGHV5-6 | CARQGGYGYSPYYFDYW | IGKV6-15 | CQQYNSYPLTF | 11 nt (6) |
| 1.30 | 1 | 29-36 | 10 | IGHV1-4 | CAVRRENYAMEYW | IGKV4-79 | CHQWSSFPPTF | 24 nt (15) |
| 1.31 | 1 | 29-36 | 10 | IGHV7-3 | CARDIPYAMDYW | IGKV1-135 | CWQGTTHFPRTF | 5 nt (0) |
| 1.33 | 1 | 29-36 | 10 | IGHV1-4 | CARGFGLYFDYW | IGKV4-57 | CHORSSYPPTF | 28 nt (19) |
| 1.37 | 1 | 37-39 | 9 | IGHV1-5 | CARTGTHWYFDVW | IGKV3-5 | CQQSNEDPWTF | 19 nt (10) |
| 1.38 | 1 | 37-39 | 9 | IGHV14-3 | CVRFGAVPRFSYW | IGKV1-117 | CFQGSHPVPTF | 18 nt (9) |
| 1.39 | 1 | 37-39 | 9 | IGHV5-6 | CAKDGGYGYGWFAYW | IGKV4-61 | CQQYHSPPTF | 14 nt (10) |
| 1.41 | 1 | 40-46 | 8 | IGHV14-1 | CCFFYGYGAWFGYW | IGKV6-17 | CQQHYTSPRTF | 19 nt (11) |
| 1.44 | 1 | 40-46 | 8 | IGHV9-3 | CARRGGNYYVYAVDYW | IGKV14-111 | CLQYDELYTF | 8 nt (5) |
| 1.47 | 1 | 47-52 | 7 | IGHV5-17 | CARQNWYDYM | IGKV12-44 | CQHHYGTPLTF | 18 nt (8) |
| 1.49 | 1 | 47-52 | 7 | IGHV1-80 | CARSGYGVFAYW | IGKV4-55 | CQQWSSYPPTF | 32 nt (13) |
| 1.50 | 1 | 47-52 | 7 | IGHV1-18 | CGRGGYGNPFAYW | IGKV4-63 | CFQGSGLPLTF | 23 nt (16) |
| 1.51 | 1 | 47-52 | 7 | IGHV9-3 | CARSPTWFAYW | IGKV15-103 | CQQGGSFPYTF | 12 nt (9) |
| 1.54 | 1 | 53-63 | 6 | IGHV1-4 | CARKSNLFPYW | IGKV6-20 | CGQTYSPPTF | 16 nt (6) |
| 1.56 | 1 | 53-63 | 6 | IGHV1-26 | CGRSYGYSYAMDYW | IGKV16-104 | CQQHNEYPTF | 25 nt (15) |
| 1.57 | 1 | 53-63 | 6 | IGHV5-17 | CARSYGSSLDYW | IGKV12-41 | CQHFWSPTPTF | 19 nt (8) |
| M1_low2 | 1 | 97-137 | 4 | IGHV1-4 | CARDYDGFAYW | IGKV1-110 | CQQTSSPSTF | 21 nt (13) |
| M1_low3 | 1 | 97-137 | 4 | IGHV1-18 | CARSGYATAFAYW | IGKV4-55 | CQQGNTLLMYTF | 26 nt (13) |
| M1_low5 | 1 | 190-248 | 2 | IGHV1-26 | CARYGNYFDYW | IGKV10-96 | CLQYASSPLTF | 19 nt (12) |
| M1_low6 | 1 | 249-404 | 1 | IGHV1-19 | CARRGDSYQFPYAMDYW | IGKV1-110 | CQQRSSYPWTF | 9 nt (6) |
| M1_low7 | 1 | 249-404 | 1 | IGHV1-26 | CARSTAIYGMIDYW | IGKV9-120 | CHQRSSYPSTF | 30 nt (15) |
| M1_low9 | 1 | 249-404 | 1 | IGHV8-11 | CARIEGLLAWFGYW | IGKV1-110 | CSQSTHVPWTF | 22 nt (8) |
| M1_low10 | 1 | 249-404 | 1 | IGHV1-82 | CARAGYGTGFAYW | IGKV4-55 | CQQWSTYPWTF | 14 nt (8) |
| M1_low12 | 1 | 249-404 | 1 | IGHV1-4 | CVRSDGDYGYVPRWFGYW | IGKV8-27 | CHQYLSSYTF | 29 nt (17) |
| M1_low14 | 1 | 249-404 | 1 | IGHV5-9-1 | CQRLRGYFDVW | IGKV1-110 | CSQSTHVPYTF | 20 nt (9) |
| M1_low15 | 1 | 249-404 | 1 | IGHV13-2 | CSRSFYDYVYAMDYW | IGKV9-124 | CLQYASYPWTF | 5 nt (2) |
| 2.2 | 2 | 2 | 32 | IGHV2-6-8 | CARDYGANNYAMDYW | IGKV4-57 | CQQRSSYPPTF | 14 nt (8) |
| 2.5 | 2 | 5 | 27 | IGHV7-3 | CARDMNGSIYWLVDVW | IGKV12-89 | CQNVLYSPWTF | 16 nt (8) |
| 2.10 | 2 | 10 | 18 | IGHV1-26 | CARNPYW | IGKV1-117 | CFQGSHPVPLTF | 26 nt (8) |
| 2.11 | 2 | 11 | 16 | IGHV1-19 | CASQLGSWFAYW | IGKV14-111 | CLQYDEFPTYF | 14 nt (5) |
| 2.13 | 2 | 12-13 | 15 | IGHV1-9 | CARSGGFYDPGRGYAMDYW | IGKV4-74 | CHQYYSRPTF | 32 nt (17) |
| 2.14 | 2 | 14-16 | 14 | IGHV8-12 | CARSYADFVW | IGKV4-80 | CHQWSSYPPTF | 9 nt (4) |
| 2.16 | 2 | 14-16 | 14 | IGHV1-55 | CTRLRYWYLDVW | IGKV14-111 | CLQYDEFPTYF | 19 nt (10) |
| 2.17 | 2 | 17-18 | 13 | IGHV9-3 | CATPDYFAMDYW | IGKV8-21 | CKQSYLPLWTF | 9 nt (6) |
| 2.19 | 2 | 19-21 | 12 | IGHV1-5 | CTRSEGYFYFDYW | IGKV14-111 | CLQYDEFPTYF | 19 nt (14) |
| 2.20 | 2 | 19-21 | 12 | IGHV5-6 | CTRRDFYGYAMDYW | IGKV8-28 | CQNDHSYPYTF | 13 nt (5) |
| 2.23 | 2 | 22-25 | 11 | IGHV6-6 | CTRDVFWAYW | IGKV1-135 | CWQGTTHFPOTF | 13 nt (11) |
| 2.26 | 2 | 26-28 | 10 | IGHV5-6 | CARRGGYGYGYFDYW | IGLV1 | CALWYSNHLV | 11 nt (6) |
| 2.28 | 2 | 26-28 | 10 | IGHV8-8 | CARDLYDDGTAYFDYW | IGKV8-27 | CHQYLSSRSF | 23 nt (8) |
| 2.29 | 2 | 29-32 | 9 | IGHV14-1 | CTPYDYDVSAFAYW | IGKV12-44 | CQHHGSPRTF | 13 nt (10) |
| 2.30 | 2 | 29-32 | 9 | IGHV1-39 | CARWGEIYPYAMDYW | IGKV10-96 | CQQGNTLPWTF | 25 nt (13) |
| 2.31 | 2 | 29-32 | 9 | IGHV1-5 | CTRCYGNPFYFDYW | IGKV4-74 | CHQHRSYPPTF | 28 nt (16) |
| 2.32 | 2 | 29-32 | 9 | IGHV2-2 | CARKGPQLVFDYW | IGKV6-23 | CQQYSSYPYTF | 19 nt (6) |
| 2.35 | 2 | 33-39 | 8 | IGHV1-81 | CARLLYGYGYLDYW | IGKV14-111 | CLQYDEFWTF | 21 nt (12) |
| 2.36 | 2 | 33-39 | 8 | IGHV1-39 | CARWGEIYPYAMDYW | IGKV10-96 | CQQGNTLPWTF | 29 nt (17) |
| 2.37 | 2 | 33-39 | 8 | IGHV1-54 | CARSDGNEDYW | IGKV4-55 | CQWSSYPFAF | 13 nt (3) |
| 2.38 | 2 | 33-39 | 8 | IGHV8-11 | CARIYYGSSSRHFDYW | IGKV3-4 | CQQSIEDPTF | 18 nt (10) |
| 2.42 | 2 | 40-48 | 7 | IGHV1-4 | CARSNDGGFAYW | IGKV4-59 | CQWSSNPPTF | 9 nt (4) |
| 2.43 | 2 | 40-48 | 7 | IGHV1-61 | CARSDDGYWFAW | IGKV3-7 | CQHSWEIPPTF | 25 nt (13) |
| 2.44 | 2 | 40-48 | 7 | IGHV1-22 | CARGLPYHGLDNW | IGKV4-61 | CQQYHSSPPTF | 27 nt (20) |
| 2.50 | 2 | 49-56 | 6 | IGHV2-2 | CAKSPYYGAMDYW | IGKV5-43 | CQQSNSWPPTF | 12 nt (7) |
| 2.52 | 2 | 49-56 | 6 | IGHV1-69 | CTRDYHGTSSMDYW | IGKV1-110 | CSQSTHVPWTF | 48 nt (19) |
| 2.53 | 2 | 49-56 | 6 | IGHV2-3 | CAKTYYGAMDYW | IGKV4-63 | CLQGSYPPTF | 22 nt (12) |
| 2.54 | 2 | 49-56 | 6 | IGHV1-26 | CARGITTVPYAMDYW | IGKV10-96 | CQQGNTLPWTF | 22 nt (12) |
| 5.8 | 5 | 8-9 | 17 | IGHV1-15 | CTREGYFYDVRWFAYW | IGKV2-109 | CAQNLELPTF | 12 nt (9) |
| 5.9 | 5 | 8-9 | 17 | IGHV14-3 | CASAATWAYWYFDVW | IGKV4-91 | CQQGSILPTF | 3 nt (2) |
| 5.10 | 5 | 10-11 | 16 | IGHV9-3 | CARGDYGNYERVAWHAYW | IGKV10-96 | CQQGDTVPPTF | 24 nt (14) |
| 5.11 | 5 | 10-11 | 16 | IGHV14-3 | CARFGNYPYWYFDVW | IGKV4-70 | CHQRSSYPYTF | 10 nt (8) |
| 5.19 | 5 | 19-21 | 11 | IGHV4-1 | CARPGFRYGYAMDYW | IGKV6-25 | CQHYSTPYTF | 10 nt (5) |
| 5.21 | 5 | 19-21 | 11 | IGHV5-9-1 | CSRDSYGYSGYGYFDVW | IGKV13-85 | CQQYWSPPPTF | 5 nt (5) |
| 5.22 | 5 | 22-24 | 10 | IGHV2-3 | CAKFSYGSYYAIDYW | IGKV6-17 | CQHYSTPLTF | 19 nt (12) |
| 5.23 | 5 | 22-24 | 10 | IGHV5-17 | CAREFAYW | IGKV12-44 | CQHHYGSPTF | 7 nt (5) |
| 5.24 | 5 | 22-24 | 10 | IGHV1-64 | CARFSGYFDVW | IGKV4-50 | CQQTSSPPTF | 20 nt (8) |
| 5.26 | 5 | 25-28 | 9 | IGHV1-18 | CARSGYDFAYW | IGKV9-124 | CLQYSTYPWTF | 33 nt (18) |
| 5.27 | 5 | 25-28 | 9 | IGHV3-1 | CARFYRSTGIAYW | IGKV4-70 | CHQRHYPWTF | 13 nt (7) |
| 5.28 | 5 | 25-28 | 9 | IGHV9-1 | CTQLGLLAWFAYW | IGKV10-94 | CQQYSKLPTF | 13 nt (6) |
| 5.30 | 5 | 29-38 | 8 | IGHV1-63 | CARKNYGSTYFDYW | IGKV1-117 | CFQGSHPWTF | 15 nt (7) |
| 5.31 | 5 | 29-38 | 8 | IGHV1-9 | CVRGRSPIYDYDDNW | IGKV17-127 | CFQSDNMPYTF | 19 nt (12) |
| 5.32 | 5 | 29-38 | 8 | IGHV1-66 | CAKNYGIHYGMDYW | IGKV13-85 | CQQYWTIPYTF | 18 nt (10) |
| 5.34 | 5 | 29-38 | 8 | IGHV5-6 | CARQVYDYGTMIDYW | IGKV10-96 | CQQGNTLPRTF | 18 nt (7) |
| 5.35 | 5 | 29-38 | 8 | IGHV8-8 | CARKAYFGDYFDYW | IGKV6-23 | CQQYSSYPLTF | 9 nt (6) |
| 5.36 | 5 | 29-38 | 8 | IGHV9-3 | CARGWGYFDYW | IGKV6-17 | CQHYSTPLTF | 19 nt (12) |
| 5.39 | 5 | 39-48 | 7 | IGHV1-4 | CARRVGEYFYFDYW | IGKV4-79 | CHQWNSYPPTF | 13 nt (8) |
| 5.40 | 5 | 39-48 | 7 | IGHV9-3 | CARAPLFYAMDYW | IGLV1 | CALWYSNHWVF | 17 nt (12) |
| 5.42 | 5 | 39-48 | 7 | IGHV1-76 | CARWGDSSGTGAMDYW | IGKV3-7 | CQHSWEIPLTF | 37 nt (12) |
| 5.43 | 5 | 39-48 | 7 | IGHV4-1 | CARPGFRYGYAMDYW | IGKV12-44 | CQHHYGTPLTF | 10 nt (5) |
| 5.45 | 5 | 39-48 | 7 | IGHV8-12 | CARLYDGYSYALDYW | IGKV12-44 | CQHHYGTPTF | 14 nt (8) |
| 5.46 | 5 | 39-48 | 7 | IGHV1-85 | CTTGDYLYVHFDVW | IGKV15-103 | CQQGQSYPTF | 30 nt (17) |
| 5.47 | 5 | 39-48 | 7 | IGHV2-9-1 | CVRGGYGYGNFYWFFDVW | IGKV1-117 | CFQGSHPVPTF | 11 nt (7) |
| 5.49 | 5 | 49-54 | 6 | IGHV1-26 | CARRGSRFYDVW | IGKV6-32 | CQQDYNPWTF | 30 nt (16) |

**Extended Data Table 2 (continued). Overview and characteristics of non-binders.** Clone ID corresponds to mouse number followed by clone index number corresponding to Fig. 1g.
| ID | Mouse | IgG clone rank in mouse | # cells | V<sub>H</sub> | CDRH3 | V<sub>L</sub> | CDRL3 | nt dist. (aa-changes) |
| --- | --- | --- | --- | --- | --- | --- | --- | --- |
| 3.3 | 3 | 2-3 | 14 | IGHV1-5 | CTRSDDYYFDYW | IGKV4-70 | CHQRSNYPLTF | 13 nt (11) |
| 3.4 | 3 | 4 | 13 | IGHV8-12 | CARTRWEKYAMDYW | IGKV19-93 | CLQYDNLTYF | 9 nt (7) |
| 4.3 | 4 | 3 | 30 | IGHV1-54 | CARSDSSGDYW | IGKV4-55 | CQQWSSYPFTF | 12 nt (5) |
| 4.5 | 4 | 5-6 | 19 | IGHV5-6 | CSRRGLVDGVYPMDYW | IGKV6-23 | CQQYRTYPTF | 21 nt (11) |
| 4.6 | 4 | 5-6 | 19 | IGHV8-12 | CVRRLLWSLDYAMDNW | IGKV10-96 | CQQGNTLPYTF | 16 nt (9) |
| 4.14 | 4 | 14 | 12 | IGHV1-19 | CAITRLDYW | IGKV4-80 | CHQWSSYPFTF | 14 nt (4) |

**Extended Data Table 3.**
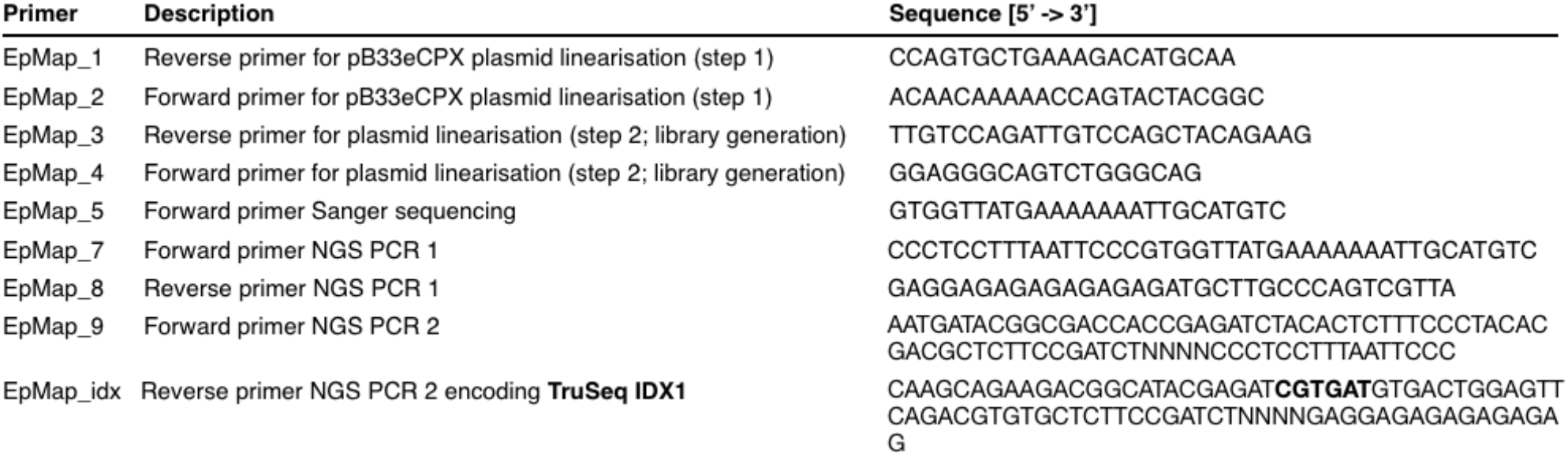
Primers used for bacterial epitope mapping and NGS library generation.

**Extended Data Table 4.**
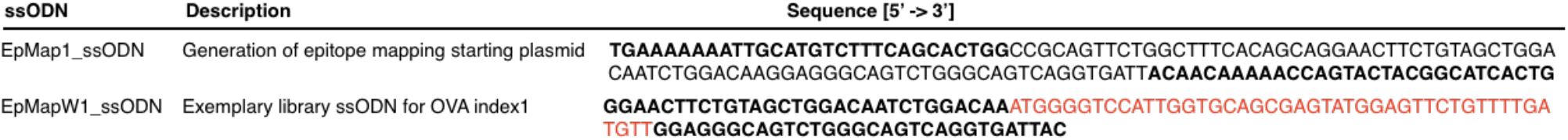
ssODNs for epitope mapping library generation. Cloning overhang sequences are indicated in bold. Insert sequence shown in red for exemplary library ssODN encodes for the first 15 a.a. of OVA.

## Notes

### Competing Interest Statement

The authors have declared no competing interest.

